# Testing for parallel genomic and epigenomic footprints of adaptation to urban life in a passerine bird

**DOI:** 10.1101/2021.02.10.430452

**Authors:** Aude E. Caizergues, Jeremy Le Luyer, Arnaud Grégoire, Marta Szulkin, Juan-Carlos Senar, Anne Charmantier, Charles Perrier

## Abstract

Identifying the molecular mechanisms involved in rapid adaptation to novel environments and determining their predictability are central questions in Evolutionary Biology and pressing issues due to rapid global changes. Complementary to genetic responses to selection, faster epigenetic variations such as modifications of DNA methylation may play a substantial role in rapid adaptation. In the context of rampant urbanization, joint examinations of genomic and epigenomic mechanisms are still lacking. Here, we investigated genomic (SNP) and epigenomic (CpG methylation) responses to urban life in a passerine bird, the Great tit (*Parus major*). To test whether urban evolution is predictable (*i.e* parallel) or involves mostly non-parallel molecular processes among cities, we analysed three distinct pairs of city and forest Great tit populations across Europe. Results reveal a polygenic response to urban life, with both many genes putatively under weak divergent selection and multiple differentially methylated regions (DMRs) between forest and city great tits. DMRs mainly overlapped transcription start sites and promotor regions, suggesting their importance in the modulation gene expression. Both genomic and epigenomic outliers were found in genomic regions enriched for genes with biological functions related to nervous system, immunity, behaviour, hormonal and stress responses. Interestingly, comparisons across the three pairs of city-forest populations suggested little parallelism in both genetic and epigenetic responses. Our results confirm, at both the genetic and epigenetic levels, hypotheses of polygenic and largely non-parallel mechanisms of rapid adaptation in new environments such as urbanized areas.

## INTRODUCTION

Identifying mechanisms involved in rapid adaptation to new environmental conditions is a central question in evolutionary biology and a is pressing task in the context of global changes of the Anthropocene ^1^. The vast majority of studies investigating mechanisms involved in rapid adaptation to new environments have focused on phenotypic plasticity on the one hand and on genetic response to selection on the other hand. At their crossroad, recent work underlines the potential role of epigenetics in rapid adaptation to new environments ^2^. In particular, environmental variations can induce differences in DNA methylation patterns and hence modulate genes’ expression and upper-level phenotypes ^3, 4^. Such methylation-linked phenotypic variations can occur during an individual’s lifetime, especially early on during the organism’s development ^5, 6^. Although methylation changes acquired across an individual’s lifetime may often be non-heritable^7^ ^but see 8, 9^, epigenetically induced phenotypic shifts may nevertheless enhance individuals fitness in new environments. Moreover, during the course of evolution, divergent genetic variants regulating epigenetic modifications may also be selected for, hence promoting the evolution of divergent epigenotypes and epigenetically-linked phenotypic variation ^10^. While epigenetic studies focused on human diseases and medical topics are now abundant, studies in an ecological context are still rare ^11^. Nevertheless, a few epigenetic studies in natural populations revealed that DNA methylation shifts might play a determinant role in local adaptation to environmental variation ^12^. There is hence an urgent need for further empirical investigations of simultaneously rapid genetic and epigenetic evolution in response to environmental change ^13^.

Urbanization rapidly and irreversibly changes natural habitats into human-made environments and is considered as a major threat to biodiversity ^14^. For species who appear to cope with urbanisation, urban habitats present a myriad of novel environmental conditions compared to the habitat where they evolved, including high levels of chemical, light and sound pollution, high proportion of impervious surfaces, high habitat fragmentation, low vegetation cover and high human densities ^15, 16^. Such extreme environmental changes compared to natural areas are expected to result in numerous new selection pressures on city-dwelling species ^17^. Accordingly, rates of recent phenotypic change, concerning multiple types of traits related to behaviour, morphology, phenology and physiology, seem to be greater in urban areas than in any other habitat types, including non-urban anthropogenic contexts ^18, 19^. The exploration of the molecular mechanisms implicated in urban-driven phenotypic changes has only begun, with both genetic ^20–22^, and epigenetic investigations ^23–25^. For instance, DNA methylation variations have been associated in vertebrates with high levels of traffic-related air pollution ^26^. Yet, epigenetic studies have been performed at relatively small genomic resolution. In addition, very little is known about the level of parallelism and hence of the predictability of genetic and epigenetic evolution in response to urbanisation in distinct cities ^27, 28^. So far, there are situations ranging from local adaptation despite strong gene flow (e.g. in the red-tailed bumblebee *Bombus lapidaries* ^29^) to restricted gene flow and independent colonization in different cities by a few founders, followed by adaptation (e.g. in the burrowing owl *Athene cunicularia,* ^30^). Providentially, recent genomic tools and virtually limitless amount of cities offer unique opportunities for comparing at high genomic resolution simultaneously individuals’ genomic and epigenomic responses in several cities to study the parallelism and predictability in molecular mechanisms implicated in rapid adaptation to urbanization ^31, 32^.

In this study, we used genome-wide and epigenome-wide sequencing approaches to compare genetic and epigenetic responses among three pairs of great tit *Parus major* populations in urban and forest habitats. At the European level, population monitoring of Great tits revealed parallel phenotypic shifts in city birds compared to their forest conspecifics, with in particular smaller and lighter urban birds laying earlier and smaller clutches ^33–36^. We investigated both average genomic (SNPs) and epigenomic (CpG methylation) differentiation and we searched for particular genomic footprints of divergent selection as well as differentially methylated regions, between forest and urban populations. Our results show that despite limited genetic differentiation and few genomic footprints of divergent selection between forest and urban populations, urban life was associated with numerous differentially methylated regions notably associated with neural development, behaviour and immunity. Hence, this study shows the potential role of epigenetic response in rapid adaptation to changing environments in urban areas. Importantly, we found little parallelism between cities in both genomic and epigenomic responses to urbanization, possibly confirming the hypothesis that multiple evolutionary ways exist to independently cope with similar novel environmental conditions.

## RESULTS

### Little genetic and epigenetic average differentiation between urban and rural populations Genetic

We used a redundancy analysis (RDA) on 74,137 SNPs obtained by RAD-sequencing to document genetic variation among the studied great tits populations, with location (Barcelona, Montpellier or Warsaw), habitat (urban vs. forest), and sex as explanatory variables. The model was highly significant (P < 0.001) but explained only a small fraction (*i.e.* less than 2%) of the total variance: R² = 0.018 (Figure 1C, SI Table S1). All three variables were significant (location: P = 0.001; habitat: P = 0.004; sex: P = 0.001). Partial RDA revealed that the net variation explained by the habitat (R² = 0.004, P = 0.001) was inferior to the net variation explained by location (R² = 0.012, P = 0.001) but higher than sex (R² = 0.002, P = 0.004, Table S1). As expected, when removing the Z chromosome from the data, sex became non-significant (P = 0.260), whereas the effects of other variables remained significant and of similar magnitude (Table S2).

**Figure 1:**
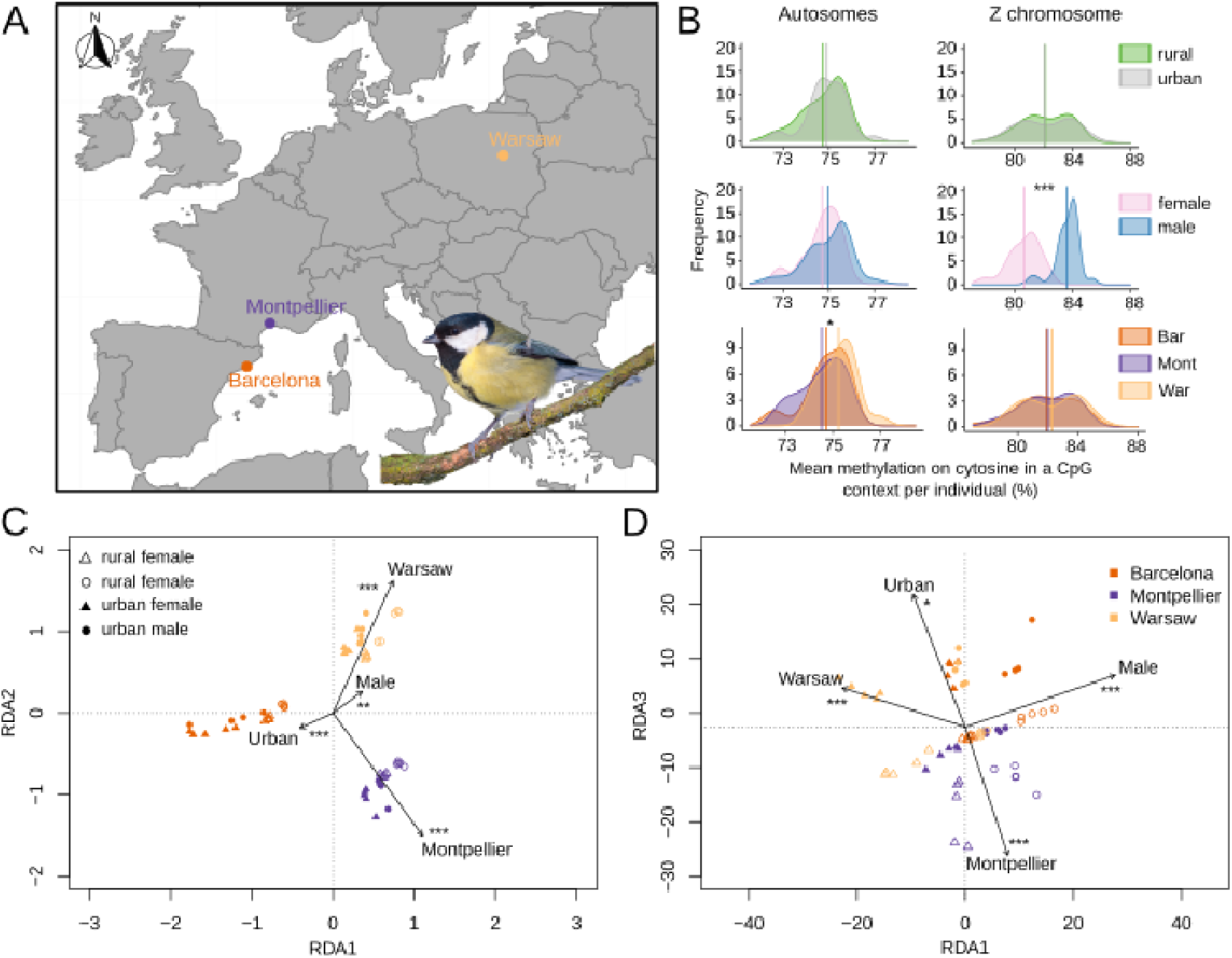
(A) Great tit blood sample locations in Europe (in urban and forest sites in and near Barcelona, Montpellier & Warsaw). (B) Distribution of mean percent of methylation on autosomes and on the Z chromosome, compared between habitats (forest versus urban), sexes and locations. (C) Redundancy analysis (RDA) on genomic data (74,137 filtered SNPs). (D) RDA on methylation levels (based on methylation levels observed at 157,741 positions). In (C) and (D), triangles represent rural habitats, circles represent urban habitats, empty and solid symbols represent females and males respectively. *** P-value < 0.001, ** P-value < 0.01 and * P-value < 0.05, related to the explanatory factors.

Genome-wide differentiation between populations, measured with Weir and Cockerham’s F_ST_ ^37^ was relatively low on average (mean F_ST_ = 0.019), in the order of 1 to 2% between habitats for each location (F_STBarcelone(rur-urb)_ = 0.018+-0.001; F_STMontpellier(rur-urb)_ = 0.012+-0.001; F_STWarsaw(rur-urb)_ = 0.018+-0.001) (see SI Table S3), suggesting relatively high gene flow and limited genetic drift among populations. Mean F_ST_ on allosomes was lower (mean F_ST_ = 0.014, Table S4) than on the Z chromosome (mean F_ST_ = 0.022, Table S5).

### Methylation

Similarly to genetic data, we performed an RDA on methylation level of 157,741 CpG sites to describe epigenetic variation among individuals in relation to location, habitat and sex. The model was significant but explained less than 1% of the total variance (R² = 0.007, P = 0.001). All variables contributed significantly (location : P = 0.001, habitat: P = 0.03, sex: P = 0.001, Table S6). Partial RDA revealed that location and sex explained a similar proportion of the total variance which was higher than habitat (location: R² = 0.003, P = 0.001; sex: R² = 0.003, P = 0.001; habitat: R² = 0.001, P = 0.3). When removing the Z chromosome from analyses, results remained similar (Table S7), showing that difference in methylation was not entirely driven by potential diverging patterns associated with sexual chromosomes.

We then investigated more finely whether individual methylation on CpG cytosines varied across location, habitat (urban vs forest), sex, and location×habitat interaction, using an ANOVA, run on autosomes and Z chromosome separately (Fig 1B, Table S8). For autosomes, we detected a significant effect of location (F = 3.319, P = 0.044), with Montpellier individuals showing lower methylation levels that Warsaw ones (Fig 1B, Tukey test: P = 0.04) and no other difference between pairs of cities. Also, no significant effect of sex (P = 0.263) or habitat (P = 0.478) was found, suggesting that urbanization did not have an important overall effect on global methylation levels. For the Z chromosome, we found a strong difference between sexes, with homogametic males showing 2.98% more methylated Z than heterogametic females (Fig 1B; P = 1.45×10-15), while no significant difference between location (P = 0.577) or habitat (P = 0.915) was found.

### Non-parallel yet strong genomic footprints of divergent selection between urban and forest populations and evidence for polygenic adaptation

We used two methods to investigate outlier SNPs potentially under divergent selection between forest and urban populations: an F_ST_-outlier based method (Bayescan^38^, v2.1, default parameters), and a multivariate method (using an RDA, ^39^) aiming respectively at identifying strong outliers indicating footprints of differentiation for each population pair and weaker outliers across the several population pairs studied at once. First, Bayescan identified 15 outliers for Barcelona, 11 for Montpellier and 10 for Warsaw, overall distributed on 15 chromosomes and associated to 13 genes in 5kb upstream or downstream regions (Figure 2, q-value < 0.1, see Figure S1). None of these outliers was shared between the three population pairs, revealing no convergence between cities. Second, using the multivariate approach based on a RDA (following the procedure for constrained RDA described earlier), we investigated the habitat effect and extracted a list of 1163 loci with outlier loading score following Forester *et al.* ^39^ method (Figure 2, see Figure S2 for threshold), suggesting a higher number of loci undergoing weaker and polygenic selection. These 1163 loci were associated with 561 genes.

**Figure 2:**
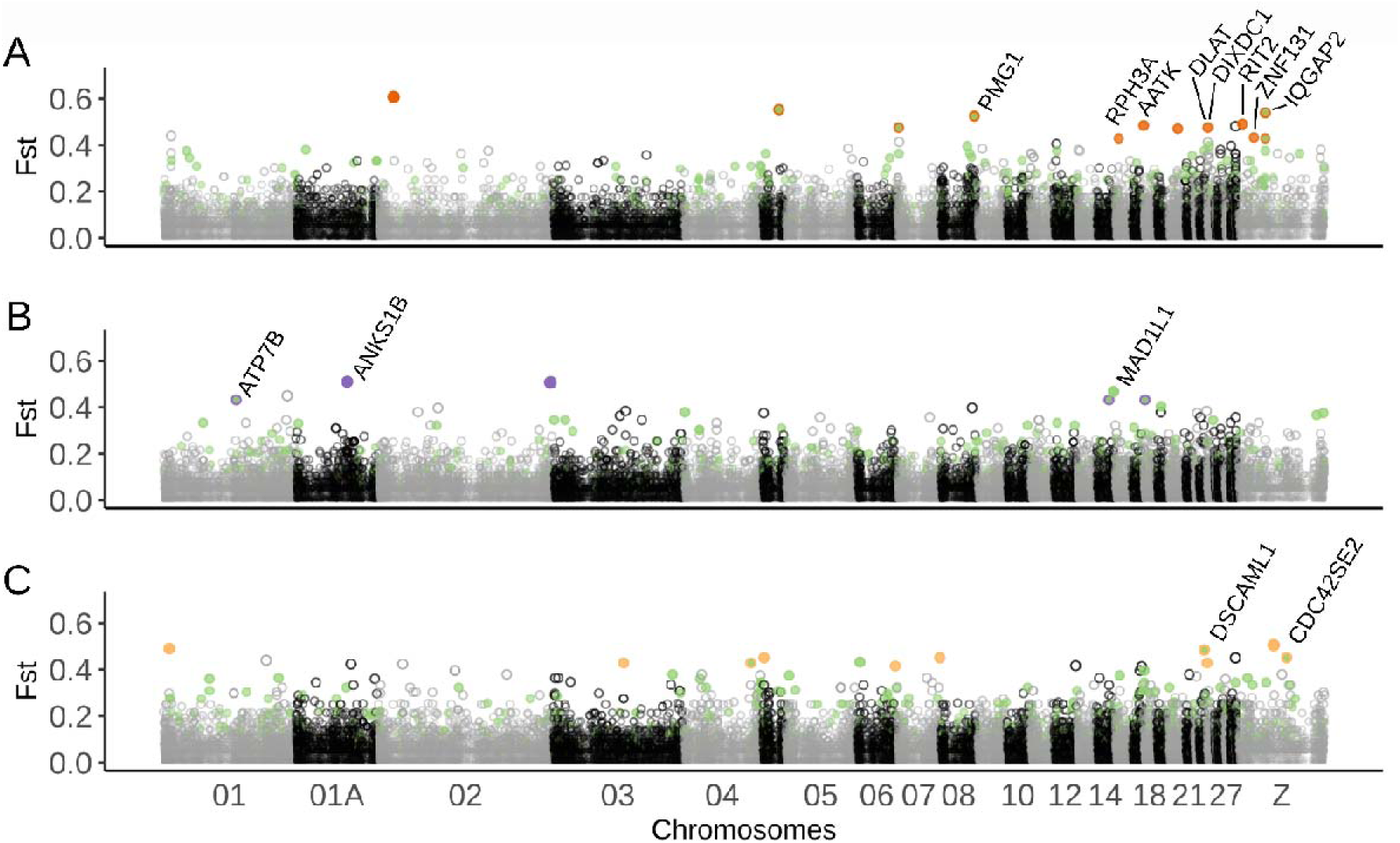
Manhattan plot of mean F_ST_ between urban and forest populations along the Great tit genome for (A) Barcelona, (B) Montpellier and (C) Warsaw. Dark orange (A), purple (B) and light orange points (C) represent significant outlier SNPs identified by the F_ST_-outlier test Bayescan for each population pair, given with their associated genes in 5 kb. Green points represent outliers found with the multivariate RDA approach. A few SNPs were identified by both the Bayescan and the RDA methods and signalled as a green point circled with the colour used for the considered pair.

To investigate the potentially statistically enriched gene ontologies (GO) for the set of genes associated to RDA outliers, we used topGO R package ^40^ with GO referenced for the chicken *Gallus gallus* (ggallus_gene_ensembl). GO analyses revealed the existence of overrepresented ontologies (P < 0.05 and at least 3 genes per GO). Among the most promising GO terms we found functions related to the nervous system (GO:0048846, axon extension involved in axon guidance; GO:0035418 protein localization to synapse; GO:0021987 cerebral cortex development; GO:0007274, neuromuscular synaptic transmission), the blood system (GO:0045777 positive regulation of blood pressure; GO:0042311, vasodilatation), hormonal response (GO:0071277, cellular response to estrogen stimuli) and stress (GO:0033555, multicellular response to stress), revealing potential important functions involved in adaptation to urban habitats. Detailed GO results are presented in SI Table S9 and Figure S3.

### Evidence for differentially methylated regions in urban environments

We identified a total of 224 distinct DMRs between urban and forest great tits: 80 for Barcelona, 68 for Montpellier and 93 for Warsaw. Only 14 DMRs (6.25%) were found repeatedly in at least two comparisons, and only 3 were common to the three cities. 7 (of these 14 parallel DMRs were in the same direction of methylation in urban compared to forest areas (Figure 3, Figure S4 A). Barcelona urban birds presented significantly more hypomethylated than hypermethylated DMRs (χ^2^ = 11.25, P < 0.001), but no difference was found for Montpellier (χ^2^ = 0.941, P = 0.332) nor for Warsaw (χ^2^ = 0.011, P = 0.917). DMRs were distributed across all the 32 chromosomes as well as on 37 unplaced scaffolds. 203 of the 224 different DMRs (91%) overlapped genes or 5kb flanking regions, among which 52% were directly located in gene bodies, promoter or TSS sequences (35.3% in gene bodies and 47.3% in promoters/TSS) suggesting their putative functional role in gene expression and regulation and/or splicing events.

**Figure 3:**
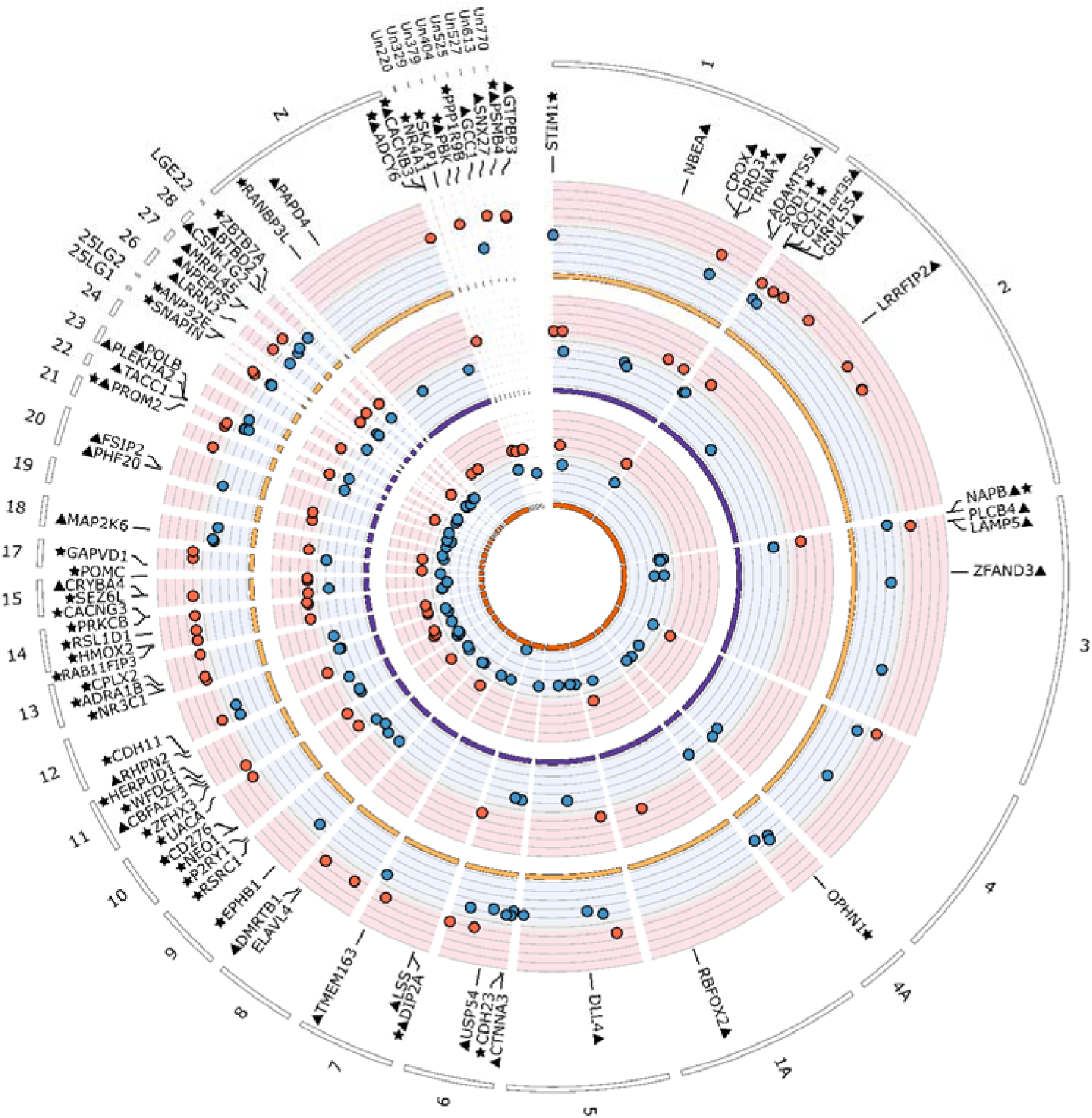
Circos plot of differentially methylated regions (DMRs) identified between populations of forest and urban great tits in and near Barcelona, Montpellier and Warsaw (from inner to outer circles). Red points show hypermethylated regions in urban great tits relatively to forest birds, and blue points show hypomethylated regions. For graphical clarity, only a subset of genes are represented: genes associated with the 10% most extreme DMR (triangles) and genes found associated with DMR in at least two cities (stars). Names of the genes found within 5 kb of the represented DMRs are given.

Following the procedure previously described, GO analyses on the pooled genes list revealed an overrepresentation of modules associated to the nervous system (GO:2000300, regulation of synaptic vesicle exocytosis; GO:0050804 modulation of synaptic transmission), immunity (GO:005728, negative regulation of inflammatory response; GO:0050852, T cell receptor signalling pathway), metabolic activity (GO:006816 calcium ion transport; GO:0055072, iron ion homeostasis, GO:0043087, regulation of GTPase activity), behaviour (GO:0007626, locomotory behavior) and endocrine processes (GO:0044060: regulation of endocrine process). All enriched GO are presented in SI Figure S5 and Table S10.

We also searched for DMRs between sexes, following the same procedure. We identified 206 DMRs associated with sex, of which 58 for Barcelona, 81 for Montpellier and 99 for Warsaw. On a total of 206 DMRs, 181 (57.3%) were on genes or in a 5kb upstream/downstream region around genes. GO analyses revealed enrichment of genes involved in development, growth and morphogenesis, among others (see detailed enriched GO Table S10 & Figure S5).

Almost twice more sex DMRs were shared between locations (11,7%) than between habitats (6.25%, see Figure S4 A & B; z-test: χ^2^-squared = 3.885, P = 0.049). When taking into account the direction of methylation difference, 7 sex DMRs were shared between at least two cities (9.7%) which was three times more than for habitat DMRs (3.1%; z-test: X-squared = 7.904, P = 0.005).

## DISCUSSION

The urban sprawl is a worldwide phenomenon deeply affecting the environment and thus requiring fast adaptive responses in city dwellers. While a large body of literature already describes a myriad of phenotypic shifts in urban populations of numerous species ^35, 41, 42^, the molecular bases of these shifts and their evolutionary implications remain yet to be documented and understood. This study uses genomic and epigenomic analyses to decipher the potential molecular bases implicated in phenotypic shifts and adaptation in several urban populations of a passerine bird, the Great tit. Note that this species shows largely parallel phenotypic shifts across its range in terms of morphology and life history (Thompson *et al., sub*). Genomic analyses revealed weak yet significant average differentiation between urban and forest populations, suggesting ongoing gene flow and limited drift in urban populations. These analyses also identified a limited number of loci putatively under strong selection, non-repeated between pairs, and numerous loci supposedly under weaker selection, compatible with a polygenic model of evolution. From the epigenomic side, we found weak average differentiation of the methylome between urban and forest birds, suggesting an absence of genome-wide epigenetic deregulations, which is notably in line with an absence of strong genetic differentiation. In turn, we identified several strongly differentially methylated regions between urban and forest birds, mostly non-repeated between pairs and hence potentially implicated with local evolution of urban populations. Genes associated with either genomic footprints of divergent selection or differentially methylated regions had relatively similar functions, related to the nervous system, metabolism, immunity and behaviour, that have been repeatedly convicted in other studies ^40^. Hence, by identifying non-repeated genetic and epigenetic responses among replicated forest-urban population pairs, our findings support the hypothesis of mostly non-parallel rapid *de novo* adaptation to similar environments via both genetic and epigenetic mechanisms. Our results are in line with accumulating evidence that polygenic adaptation and epigenetic reprogramming may be involved in quick phenotypic shifts in response to rapidly emerging constraints such as urbanization.

Overall genetic differentiation between populations was relatively low (F_ST_ ranging from 0.9% to 3.4%), although higher than what has been found at a much larger scale across the species distribution (e.g. around 1% of F_ST_ between UK and Spanish or French and Spanish populations ^41^). Low but significant differentiation levels are in line with previously documented divergences between city and forest great tit populations ^20, 45^, which altogether suggests important gene flow, large effective population sizes and limited genetic drift at multiple spatial scales ^46^. This overall genetic context is particularly suitable to search for genomic footprints of divergent selection between urban and forest populations, which would easily be identifiable above the neutral level of genetic differentiation.

We found a limited number of strong footprints of divergent selection, which is in line with previous results of Perrier et al. ^20^ in Montpellier, and of Salmon et al. ^22^ across nine European cities. Similarly to low levels of parallelism in allele frequency changes between cities observed by previous studies ^22, 47^, and despite similarities in phenotypic shifts in the European cities ^35^, none of these outliers were shared between cities. This result suggests limited parallel evolution, supporting a scenario of independent *de novo* evolution between cities and/or different selection pressures between cities. Indeed, there may be multiple evolutionary solutions to the same environmental challenges ^48^ and multiple traits are linked to the same functional outcome ^49^. Besides, the identification of numerous outliers by the multivariate framework applied at the scale of all six sampling sites supports a model of polygenic urban adaptation implicating multiple genes, biological pathways and phenotypic traits ^50^. Polygenic adaptation may be expected in urban habitats since the multiple new environmental conditions in cities most probably result in many novel selective pressures acting on a multitude of functional traits ^51^, and because many of these traits may be quantitative, and genetically correlated ^52^. Further polygenic analyses on more samples and more markers (i.e. whole genome data) are however required to increase our capacity to estimate more precisely the potential effect of each genetic variant implicated in the adaptation to life in the city ^53, 54^.

Several genomic footprints of divergent selection between urban and forest environments were in, or in the vicinity of, genes that have already been described as playing a role in neuronal development, behaviour or cognitive abilities. In particular, the NR4A2 gene plays an important role in recognition of novel objects and memory in mice ^55^. Reaction to novel objects and novel food is known as one of the main factors determining the capacity of a species to thrive in an urban environment ^41^. The DCX gene is related to neuronal plasticity ^56^ and experimental approaches revealed that artificial light at night induces an overexpression of this gene linked to a change in behaviour and expression of depressive-like behaviour in crows ^57^. Finally, the CHRNA1 gene is associated with aggressive behaviour in chicken ^58^, and higher aggressiveness is commonly observed and hypothesized as adaptive in urban bird populations ^59^. Besides, the gene ontology enrichment analysis, performed on the entire set of genes identified via the outlier genome scan, reinforced these findings since multiple enriched GO terms were associated with the nervous system and stress response as well as hormonal response. These results are informative on the type of traits involved in avian urban adaptation in cities and corroborate previous results from ^22, 60, 61^ suggesting that natural selection repeatedly acted on neuronal, behavioural and cognitive traits that could contribute to the phenotypic shifts described in urban great tits (i.e. more aggressive and exploratory birds, with higher breath rate : ^62, 63^ & Caizergues *et al. in prep*).

Contrary to the common prediction that living in cities is likely to influence epigenomes ^24, 25^, no genome wide pattern of differentiation in methylation between urban and forest great tits was detected. However, we observed a difference in mean methylation level between birds from Warsaw and Barcelona on their autosomes, as well as between males and females on the Z chromosome, showing that methylation differences were identifiable. In addition, we found no mean difference in methylation level between habitats, revealing that urbanization did not strongly affect overall methylation levels in a specific direction. This overall low differentiated methylation context is perfectly suitable to investigate more localized zones that could differ in their levels of methylation. Note that the strong methylation contrast between males and females, particularly on sex chromosomes (Figure 1B), is in line with previous reports in vertebrates ^64, 65^ showing that methylation plays a major role in sex differentiation via regulation of gene expression and genetic imprinting.

Despite the non-significant effect of habitat on overall methylation levels, we found a large number of DMRs between pairs of forest and urban populations, suggesting that urbanization did affect particular regions of the genome. DMR were less likely to occur within a gene body than by chance, but it was not the case for promoter or TSS regions. This latter result contrasts with Watson and collaborators ^25^ who recently found an under-representation of DMR in both gene body and regulatory regions in urban great tits from Malmö (Sweden). Across the three cities, 35.3% of DMR were directly localized in gene bodies, and 47.3% in TSS or promoter regions, suggesting a potential role in gene expression modulation. Direction of methylation patterns did not follow any absolute pattern (no over-representation of hypo- or hypermethylated DMR in urban birds, Figure 3), in line with Watson and collaborators’ analyses on blood sample. However, birds in Barcelona presented significantly more hypomethylated DMR than hypermethylated ones, a result that warrants further investigations.

Only a limited number of urbanization-linked DMR were shared between two or more locations. In contrast, three times more sex linked DMRs were found in two locations or more. This comparison suggests that urbanization-linked epigenetic modifications most probably do not occur in a parallel way across cities, but rather that each city might have its particular epigenetic response. Indeed, in the study field of urban evolutionary biology, cities are often regarded as valuable replicates of human induced habitats ^31, 66^, and it is often expected that parallel adaptive responses to similar selective pressures will occur. This expectation is particularly strong when phenotypes show parallel changes, as is the case for the Great tit, which is consistently smaller and lays earlier and smaller broods in the various cities where it has been studied, compared to forest habitats ^67, 68^. However, as discussed above, parallel adaptation to similar environmental conditions should not be expected in the case of independent evolution, especially for multilocus traits. Additionally, cities are different from each other because of a wide array of climatic, cultural, historical and socioeconomic factors ^15^. In fact, besides the obvious differences in cities’ climatic conditions depending on their position on the globe, land use, fragmentation and pollutants levels can also largely vary across cities ^69^. In a general way, pollutants are known to affect DNA methylation and result in both hypo and hypermethylation, but the patterns of change observed largely rely on the pollutant involved ^70^. In this case, differences in cohorts of pollutants present in cities could be responsible for differences in patterns of methylations. Taken together, these results highlight the importance of questioning the assumption that cities are replicated environments that can be considered similar.

As mentioned earlier, increasing evidence suggests that DNA methylation can be associated with environmental and stress factors (env: ^12^, stress: ^71^) especially during early development ^72^. Here, we found four genes (POMC, ADAMTS3, PAPD4 et GCC1), associated with DMR that were previously described in great tits as undergoing major changes in methylation levels in case of exposure to higher levels of pollutants ^73^. Notably, the functions of these genes remain to be determined, and they could thus be interesting to target in future studies. In addition, the past literature has repeatedly found SERT and DRD4 as two major genes involved in urban specific avian human avoidance (or wariness) behaviours (see for example Riyahi *et al.* ^23^ in the Great tit, Garroway and Sheldon ^74^ in the blackbird, Van Dongen *et al.* ^75^ in the black swan & Mueller *et al.* ^76^ in the burrowing owl) (see SI Figure S8 to S12 to see patterns of methylation associated with 6 classical great tit linked candidate genes). In this study, while urban great tits show higher levels of aggressiveness in at least two of the cities (^43^ & Caizergues *et al. in prep*) we found no DMR associated with these two genes in either of the three forest-city pairs. However, we found a significant urban-related change in methylation linked to the DRD3 gene, belonging to the same gene family as DRD4 and known to be similarly involved in chicken aggressive behaviour ^77^. In line with these results, GO analyses revealed enrichment in genes associated with neuronal functions, behaviour, but also blood, immune and endocrine systems (Table S9, Figure S3), revealing the potential need of physiological adjustments in urban habitats. Surprisingly, a recent study on great tit differences of methylation between city and forest habitats in another European city found no GO enrichment in blood, while some in liver tissue^25^ (note that they investigated DMSs, Differentially Methylated Sites, which differs from DMRs identified here). These contrasted results highlight the fact that methylation patterns highly depend on the analysed tissues ^11^, and show, once more, that urban linked methylation might not be similar from one city to another. In addition, it has been demonstrated that DNA methylation can undergo seasonal variation ^78^. Hence, analyses on multiple tissues and life-stages replicated in multiple pairs of urban and forest populations will allow to draw a broader view of the impact of urbanization on global methylation patterns and to understand replicated parallel occurrence across cities. However, tissue-specific and age-specific analyses in multiple individuals across several pairs of urban and forest environments poses major technical, budget and ethical limitations and should be coordinated very carefully. Additionally, specific drivers of shifts in methylation remain to be disentangled to understand which environmental factors are responsible for which change in methylation. To do so, experimental settings manipulating environmental factors such as performed by Mäkinen and collaborators ^73^ would be particularly useful. More integratively, information on how shifts in methylation patterns affect phenotype, fitness, and adaptation, often remain elusive ^79^. To our knowledge, a limited number of studies attempted to link methylation and expression levels in natural population contexts ^11, 44^, and even fewer in urban habitats (but see e.g. ^24, 25^). Hence, future work might need to tackle the question of the origin and adaptive significance of these variations in a controlled framework.

## CONCLUSION

In this study, we found genomic footprints of selection and differentially methylated regions associated with urbanization, suggesting that genomic as well as epigenetic processes could play an important role in the rapid adaptation to the urban habitat. To our knowledge, our study is the first to use replicated pairs of cities and forest habitats when studying urban linked methylation, offering a great opportunity to investigate convergent responses to anthropogenic environmental conditions. Taken together, our results revealed limited parallelism between cities regarding selection as well as methylation patterns, suggesting that cities might not present exactly similar environmental conditions or that different genetic pathways are involved in adaptation to urban environmental conditions, while still associated with similar biological functions. Furthermore, we highlight the need to unravel both environmental origins and evolutionary implications of methylation shifts, to understand to which extent environmental induced methylation can contribute to adaptation.

## MATERIAL AND METHODS

### Study sites and sampling

Three pairs of great tit populations in urban and forest environments were sampled in the three European cities of Barcelona (Spain), Montpellier (France) and Warsaw (Poland). For each location, 10 individuals were sampled within the city and 10 individuals were sampled from nearby forest. Blood samples were collected from breeding individuals during spring between 2016 and 2018 (except 2 individuals for Barcelona city collected in 2014 and 2015) and kept in alcohol or queen’s buffer. Samples had balanced sex ratio (5 males and 5 females for each population) except for the forest population of Barcelona where 6 females and 4 males were sampled.

### DNA extraction, RAD-seq and Reduced-Representation Bisulfite Sequencing

We used QIAGEN DNeasy blood and tissue kit to extract genomic DNA from blood samples following the provided instructions for nucleated blood samples. DNA was quantified using a NanoDrop ND8000 spectrophotometer and a Qubit 2.0 fluorometer with the DNA HS assay kit (Life Technologies). DNA quality was examined on agarose gels. We then performed RAD-sequencing and RRBS-sequencing using standard protocols. For RAD-sequencing (restriction-site-associated DNA sequencing^80^), the library preparation was done by the Montpellier GenomiX (MGX) platform (CNRS, Montpellier), using the enzyme SbfI. Each individual was identified using a unique six nucleotides tag, individuals were randomly multiplexed in equimolar proportions by libraries of 37 individuals. Each library was sequenced on a lane of an Illumina HiSeq 2500. Paired-end sequencing was used to produce 150 bp reads. This generated an average of 4.9M reads per individual. The DNA of 60 individuals were processed twice to test for reliability of the genotyping process. The RRBS-sequencing started with DNA digestion using MspI restriction enzymes, which cuts CCGG sites and targets regions that are CG rich, permitting to have a high proportion sequences in promoter regions. Each individual was identified using a unique six nucleotides tag. Individuals were randomly multiplexed in equimolar proportions by libraries of 10 individuals. Bisulfite treatment hence converted unmethylated cytosines into uracil, then converted to thymine after PCR amplification. Each library was then sequenced on a lane of an Illumina HiSeq 2500. Paired-end sequencing was used to produce 50 bp reads. This generated an average of 19.3M reads per individual.

### Genomic analyses

Fastp v. 0.19.7 ^81^ was used to trim the RAD-seq reads, keeping reads with a minimum quality of 15 before mapping individual sequences against the reference genome of the Great tit (^44^, GenBank assembly accession: GCA_001522545.3) with BWA v0.7.17 ^82^. Genotyping was conducted with stacks v2.41 ^83^ “*gstacks*” and “*population*” functions, using “*snp*” model, filtering for mapping quality > 10, alpha = 0.05, minor allele frequency > 0.05 and observed heterozygosity < 0.65. Loci were retained if present in at least 90% of individuals in each population. Loci with extreme low or high coverage were removed (5% extremes filtered out using vcftools v0.1.15 ^84^). After filtering, 74,137 SNPs were retained for subsequent population genomic analyses.

To document genomic variation among urban and rural great tits from the three locations we used a redundancy analyses (RDA), with location (Barcelona, Montpellier and Warsaw), environment (urban or rural) and sex as explanatory variables. Partial RDA was also produced to test for each variable effect (environment, location or sex) alone after controlling for all other variables. The effect of a given factor was considered significant with a p-value < 0.05.

To estimate genome-wide differentiation between populations we used Weir and Cockerham’s F_ST_ ^37^ computed using the StAMPP R package ^85^. Average F_ST_ was estimated using all SNPs, and confidence intervals were assessed using 1000 bootstrap replicates.

To identify SNPs potentially under divergent selection between urban and forest habitats we first used Bayescan v2.1 ^38^. Analyses were run for each pair of populations (Barcelona, Montpellier & Warsaw) separately, with default parameter option. As recommended by ^38^ we considered SNPs as outliers when they displayed a q-value above the 0.1 threshold.

To detect weaker footprints of divergent selection typically expected in polygenic adaptations in response to complex environmental heterogeneity, a multivariate method was used ^39^. Following a similar procedure as described above for the RDA analysis, we used a constrained RDA to investigate the effect of habitat (forest versus city) and to identify outliers SNPs displaying more than3 times SD from the mean score on the constrained axis.

### Methylation calling and analyses

The RRBS reads were first trimmed using fastp software v0.19.7 ^81^, and quality filtered keeping only reads with a quality > 15. BISMARK software v0.20.0 ^86^ was used for mapping reads on the masked reference genome with default parameters and 1 maximum allowed mismatches (-N 1). Methylation information for cytosine in a CpG context with sufficient coverage (≥10x) was extracted. An anova on mean individual methylation was run to test for location, habitat and sex effect on overall methylation levels.

Similarly, to genetic differentiation, an RDA was performed to describe epigenomic variation in function of location, habitat and sex. A partial RDA was also conducted to test for the habitat effect alone.

To identify differentially methylated regions (DMR) logistic regression including sex as covariate was performed after tiling regions of 1000pb for each location using MethylKit R package *“calculateDiffMeth”* function. Only DMR having > 10% overall difference between forest and urban habitat and q-value < 0.001 were kept. We then investigated if DMRs overlapped genes in 5 kb upstream and downstream regions using bedtools.

### Genes associated to genomic outliers and DMRs, and gene ontology analyses

We investigated if genomic outliers and DMR overlapped genes in 5 kb upstream and downstream regions using BEDTools v2.28.0 ^87^. Gene ontology analyses were performed with the R package topGO ^40^ to identify potential statistically enriched to investigate if on the list of all genes associated to RDA genomic outliers and the list of all genes associated to DMR.

## ACKNOWLEDGMENTS

We thank all the people that contributed to fieldwork and blood sample collection, in particular the members of the PLT platform of the CEFE, as well as the fieldwork teams of Barcelona and Warsaw. We also thank Patricia Sourouille and the CEFE GEMEX platform for laboratory support, and Enrique Ortega-Aboud as well as Rémi Allio for their bioinformatics support. This research was funded by the European Research Council (Starting grant ERC-2013-StG-337365-SHE to AC) and the OSU-OREME, the Polish National Science Centre (Grants Sonata BIS 2014/14/E/NZ8/00386 and Opus 2016/21/B/NZ8/03082 to MS), and the Ministry of Economy and Competitivity, Spanish Research Council (CGL-2016-79568-C3-3-P).

## SUPPLEMENTARY INFORMATION

### SUPPLEMENTARY TABLES

**Table S1:**
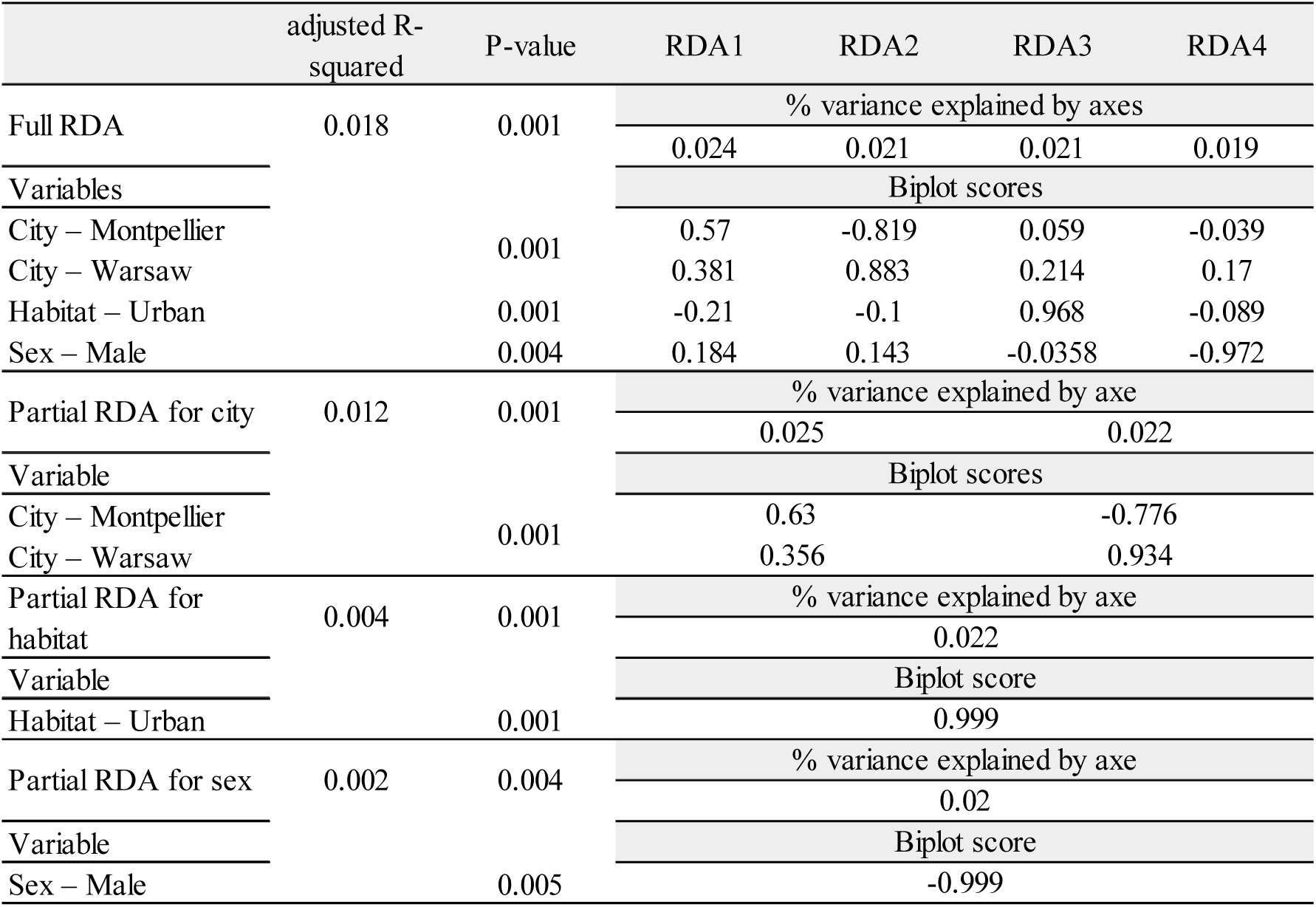
Redundancy analysis (RDA) performed on the genetic data including the Z chromosome.

**Table S2:** Redundancy analysis (RDA) performed on the genetic data without Z chromosome.

|  | adjusted R-squared | P-value | RDA1 | RDA2 | RDA3 | RDA4 |
| --- | --- | --- | --- | --- | --- | --- |
| Full RDA | 0.016 | 0.001 | % variance explained by axes |  |  |  |
|  |  |  | 0.024 | 0.021 | 0.021 | 0.017 |
| Variables |  |  | Biplot scores |  |  |  |
| City – Montpellier |  | 0.001 | 0.5725 | -0.818 | 0.049 | -0.0244 |
| City – Warsaw |  |  | 0.382 | 0.889 | 0.227 | 0.11 |
| Habitat – Urban |  | 0.001 | -0.215 | -0.119 | 0.967 | -0.073 |
| Sex – Male |  | 0.26 | 0.14 | 0.096 | -0.033 | -0.985 |
| Partial RDA for city | 0.012 | 0.001 | % variance explained by axe |  |  |  |
|  |  |  | 0.025 |  | 0.022 |  |
| Variable |  |  | Biplot scores |  |  |  |
| City – Montpellier |  | 0.001 | 0.624 |  | -0.781 |  |
| City – Warsaw |  |  | 0.362 |  | 0.932 |  |
| Partial RDA for habitat | 0.004 | 0.001 | % variance explained by axe |  |  |  |
|  |  |  | 0.022 |  |  |  |
| Variable |  |  | Biplot score |  |  |  |
| Habitat – Urban |  | 0.001 | 0.999 |  |  |  |
| Partial RDA for sex | 0.0004 | 0.373 | % variance explained by axe |  |  |  |
|  |  |  | 0.018 |  |  |  |
| Variable |  |  | Biplot score |  |  |  |
| – Urban |  | 0.369 | -0.999 |  |  |  |

**Table S3:** Fst estimation between pairs of subpopulations. 95% confidence computed intervals in brackets were computed using StAMPP package with 1000 bootstrap.

|  | Mont-Rur | Mont-Urb | Var-Rur | Var-Urb | Bar-Rur |
| --- | --- | --- | --- | --- | --- |
| Mont-Urb | 0.012 (0.011 – 0.013) | - | - | - | - |
| Var-Rur | 0.009 (0.009 – 0.010) | 0.018 (0.017 – 0.019) | - | - | - |
| Var-Urb | 0.018 (0.017 – 0.018) | 0.026 (0.025 – 0.027) | 0.018 (0.017 – 0.019) | - | - |
| Bar-Rur | 0.009 (0.009 – 0.010) | 0.019 (0.017 – 0.018) | 0.009 (0.009 – 0.010) | 0.018 (0.017 – 0.019) | - |
| Bar-Urb | 0.025 (0.024 – 0.027) | 0.034 (0.033 – 0.035) | 0.025 (0.025 – 0.026) | 0.034 (0.033 – 0.034) | 0.018 (0.017 – 0.019) |

**Table S4:** Fst estimation averaged on autosomes between pairs of subpopulations. 95% confidence computed intervals in brackets were computed using StAMPP package with 1000 bootstrap.

|  | Mont-Rur | Mont-Urb | Var-Rur | Var-Urb | Bar-Rur |
| --- | --- | --- | --- | --- | --- |
| Mont-Urb | 0.017 (0.017 – 0.017) | - | - | - | - |
| Var-Rur | 0.018 (0.018 – 0.018) | 0.015 (0.014 – 0.015) | - | - | - |
| Var-Urb | 0.014 (0.014 – 0.015) | 0.011 (0.011 – 0.011) | 0.015 (0.014 – 0.015) | - | - |
| Bar-Rur | 0.018 (0.018 – 0.018) | 0.015 (0.014 – 0.015) | 0.018 (0.018 – 0.018) | 0.014 (0.014 – 0.014) | - |
| Bar-Urb | 0.011 (0.011 – 0.011) | 0.008 (0.007 – 0.008) | 0.011 (0.011 – 0.011) | 0.008 (0.007 – 0.008) | 0.014 (0.014 – 0.014) |

**Table S5:** Fst estimation averaged on Z chromosome between pairs of subpopulations. 95% confidence computed intervals in parenthesis were computed using StAMPP package with 1000 bootstrap.

|  | Mont-Rur | Mont-Urb | Var-Rur | Var-Urb | Bar-Rur |
| --- | --- | --- | --- | --- | --- |
| Mont-Urb | 0.013 (0.006 – 0.018) | - | - | - | - |
| Var-Rur | 0.016 (0.009 – 0.021) | 0.020 (0.013 – 0.025) | - | - | - |
| Var-Urb | 0.025 (0.019 – 0.031) | 0.025 (0.020 – 0.032) | 0.020 (0.015 – 0.025) | - | - |
| Bar-Rur | 0.015 (0.010 – 0.019) | 0.017 (0.011 – 0.022) | 0.014 (0.009 – 0.019) | 0.018 (0.015 – 0.024) | - |
| Bar-Urb | 0.031 (0.026 – 0.036) | 0.041 (0.032 – 0.047) | 0.028 (0.022 – 0.034) | 0.036 (0.030 – 0.043) | 0.014 (0.009 – 0.021) |

**Table S6:** Redundancy analysis (RDA) performed on the methylation data including the Z chromosome.

|  | adjusted R-squared | P-value | RDA1 | RDA2 | RDA3 | RDA4 |
| --- | --- | --- | --- | --- | --- | --- |
| Full RDA | 0.007 | 0.001 | % variance explained by axes |  |  |  |
|  |  |  | 0.021 | 0.019 | 0.018 | 0.016 |
| Variables |  |  | Biplot scores |  |  |  |
| City – Montpellier |  | 0.001 | 0.21 | 0.259 | -0.648 | 0.685 |
| City – Warsaw |  |  | -0.603 | -0.77 | 0.198 | -0.067 |
| Habitat – Urban |  | 0.03 | -0.256 | 0.286 | 0.669 | 0.637 |
| Sex – Male |  | 0.001 | 0.736 | -0.562 | 0.263 | 0.269 |
| Partial RDA for city | 0.003 | 0.001 | % variance explained by axe |  |  |  |
|  |  |  | 0.021 |  | 0.018 |  |
| Variable |  |  | Biplot scores |  |  |  |
| City – Montpellier |  | 0.001 | -0.356 |  | 0.934 |  |
| City – Warsaw |  |  | 0.986 |  | -0.161 |  |
| Partial RDA for habitat | 0.001 | 0.3 | % variance explained by axe |  |  |  |
|  |  |  | 0.019 |  |  |  |
| Variable |  |  | Biplot score |  |  |  |
| Habitat – Urban |  | 0.181 | 0.999 |  |  |  |
| Partial RDA for sex | 0.003 | 0.001 | % variance explained by axe |  |  |  |
|  |  |  | 0.021 |  |  |  |
| Variable |  |  | Biplot score |  |  |  |
| Sex – Male |  | 0.001 | 0.999 |  |  |  |

**Table S7:** Redundancy analysis (RDA) performed on the methylation data without Z chromosome.

|  | adjusted R-squared | P-value | RDA1 | RDA2 | RDA3 | RDA4 |
| --- | --- | --- | --- | --- | --- | --- |
| Full RDA | 0.006 | 0.001 | % variance explained by axes |  |  |  |
|  |  |  | 0.021 | 0.019 | 0.018 | 0.016 |
| Variables |  |  | Biplot scores |  |  |  |
| City – Montpellier |  | 0.001 | 0.231 | -0.204 | -0.666 | 0.679 |
| City – Warsaw |  |  | -0.645 | 0.761 | 0.2467 | -0.058 |
| Habitat – Urban |  | 0.019 | -0.261 | -0.365 | 0.632 | 0.632 |
| Sex – Male |  | 0.001 | 0.698 | 0.569 | 0.326 | 0.289 |
| Partial RDA for city | 0.003 | 0.001 | % variance explained by axe |  |  |  |
|  |  |  | 0.02 |  | 0.018 |  |
| Variable |  |  | Biplot scores |  |  |  |
| City – Montpellier |  | 0.001 | -0.357 |  | 0.933 |  |
| City – Warsaw |  |  | 0.987 |  | -0.159 |  |
| Partial RDA for habitat | 0.001 | 0.176 | % variance explained by axe |  |  |  |
|  |  |  | 0.019 |  |  |  |
| Variable |  |  | Biplot score |  |  |  |
| Habitat – Urban |  | 0.168 | 0.999 |  |  |  |
| Partial RDA for sex | 0.003 | 0.001 | % variance explained by axe |  |  |  |
|  |  |  | 0.021 |  |  |  |
| Variable |  |  | Biplot score |  |  |  |
| Sex – Male |  | 0.001 | 0.999 |  |  |  |

**Table S8:** Results of Type I ANOVA performed on mean methylation level of cytosines in a CpG context per individual, including: habitat (rural vs urban), city (Barcelona, Montpellier and Warsaw) and sex as explanatory variables. The first part of the table shows results of the analysis on autosomes and the second part shows results of the analysis on the Z chromosome only.

| Autosomes |  |  |  |  |  |
| --- | --- | --- | --- | --- | --- |
| Effect | Df | Sum Sq | Mean Sq | F value | P |
| Habitat | 1 | 0.416 | 0.416 | 0.511 | 0.478 |
| Sex | 1 | 1.041 | 1.041 | 1.278 | 0.263 |
| City | 2 | 5.408 | 2.704 | 3.319 | 0.044 |
| Habitat* City | 2 | 0.316 | 0.158 | 0.194 | 0.825 |
| Residuals | 53 | 42.174 | 0.8146 |  |  |
| Z chromosome |  |  |  |  |  |
| Effect | Df | Sum Sq | Mean Sq | F value | P |
| Habitat | 1 | 0.012 | 0.012 | 0.011 | 0.915 |
| Sex | 1 | 133.584 | 133.584 | 125.077 | 1.45x10-15 |
| City | 2 | 0.979 | 0.489 | 0.458 | 0.635 |
| Habitat* City | 2 | 1.1879 | 0.594 | 0.556 | 0.577 |
| Residuals | 53 | 56.605 | 1.068 |  |  |

**Table S9:** Significantly enriched GO associated with genes overlapping 5kb windows around genomic outliers between forest and urban habitats.

| GO.ID | Term | Annotated | Significant | Expected | Weight | Associated Genes |
| --- | --- | --- | --- | --- | --- | --- |
| GO:2000618 | regulation of histone H4-K16 acetylation | 5 | 3 | 0.34 | 0.003 | ATG5, AUTS2, SIRT1 |
| GO:1902287 | semaphorin-plexin signaling pathway invo... | 7 | 3 | 0.48 | 0.009 | PLXNA1, PLXNA2, PLXNA4 |
| GO:1901842 | negative regulation of high voltage-gate... | 3 | 3 | 0.21 | 0.000 | DYSF, GEM, RRAD |
| GO:0090335 | regulation of brown fat cell differentia... | 10 | 4 | 0.69 | 0.005 | FTO, PIM1, PRDM16, SIRT1 |
| GO:0071391 | cellular response to estrogen stimulus | 6 | 3 | 0.41 | 0.006 | AR, CRHBP, ESR1 |
| GO:0071277 | cellular response to calcium ion | 28 | 7 | 1.93 | 0.002 | CRHBP, ECT2, HPCA, RASAL1, SYT1, SYT11, SYT9 |
| GO:0050953 | sensory perception of light stimulus | 57 | 6 | 3.92 | 0.026 | CDH23, CRYBB2, NYX, PCDH15, RPE65, WHRN |
| GO:0050765 | negative regulation of phagocytosis | 7 | 3 | 0.48 | 0.009 | ATG5, DYSF, SYT11 |
| GO:0048846 | axon extension involved in axon guidance | 23 | 4 | 1.58 | 0.026 | NRP2, PLXNA4, SEMA4G, SLIT3 |
| GO:0048026 | positive regulation of mRNA splicing vi... | 6 | 3 | 0.41 | 0.006 | HMX2, NCBP1, NUP98 |
| GO:0045777 | positive regulation of blood pressure | 9 | 3 | 0.62 | 0.026 | ADRA1D, GRIP2, WNK1 |
| GO:0044351 | macropinocytosis | 6 | 3 | 0.41 | 0.006 | DOCK2, MAPKAPK2, SNX33 |
| GO:0043410 | positive regulation of MAPK cascade | 197 | 15 | 13.56 | 0.013 | ADRA1D, ADRB2, APP, AR, AXIN1, EPHA8, FGFR3, MTURN, NRG1, PDGFRB, PLCB1, RET, ROR1, ROR2, SH3RF2 |
| GO:0043087 | regulation of GTPase activity | 194 | 17 | 13.36 | 0.025 | AGAP3, ARHGAP17, ARHGAP45, ARHGEF12, DOCK2, ECT2, IQGAP2, MLST8, PLCB1, PLXNA1, PLXNA2, PLXNA4, RASAL1, RDX, RGS8, SIPA1L1, WNK1 |
| GO:0042535 | positive regulation of tumor necrosis fa... | 6 | 3 | 0.41 | 0.006 | APP, HSPB1, MAPKAPK2 |
| GO:0042311 | vasodilation | 13 | 4 | 0.89 | 0.010 | ADRB2, ATG5, CPS1, NOS1 |
| GO:0035418 | protein localization to synapse | 26 | 5 | 1.79 | 0.009 | BSN, GPC4, GRIP2, HSPB1, NRXN1 |
| GO:0033555 | multicellular organismal response to str... | 37 | 6 | 2.55 | 0.006 | BRINP1, DBH, NOS1, NR4A2, PPP3CA, RET |
| GO:0021987 | cerebral cortex development | 56 | 8 | 3.86 | 0.013 | CCDC85C, CNTNAP2, DCX, HIF1A, NRP2, PAX5, PLCB1, TRAPPC9 |
| GO:0016339 | calcium-dependent cell-cell adhesion via... | 15 | 4 | 1.03 | 0.016 | CDH13, CDH17, CDH19, CDH20 |
| GO:0009755 | hormone-mediated signaling pathway | 87 | 12 | 5.99 | 0.014 | AR, ASIP, CRHBP, ESR1, ESRRB, KMT2D, NR5A1, PPARG, PRLR, REN, RNF14, SIRT1 |
| GO:0008344 | adult locomotory behavior | 46 | 7 | 3.17 | 0.006 | APP, CHL1, FGF12, NR4A2, NTAN1, PCDH15, SEZ6L |
| GO:0007274 | neuromuscular synaptic transmission | 9 | 4 | 0.62 | 0.002 | ADARB1, CHRNA1, FCHSD2, SLC5A7 |
| GO:0007171 | activation of transmembrane receptor pro... | 8 | 3 | 0.55 | 0.014 | DGKQ, NRG1, PRLR |
| GO:0007156 | homophilic cell adhesion via plasma memb... | 57 | 11 | 3.92 | 0.001 | CDH13, CDH17, CDH19, CDH20, CDH23, CNTN4, DSCAML1, IGSF11, MYOT, PCDH15, RET |
| GO:0007155 | cell adhesion | 508 | 53 | 34.97 | 0.015 | ADAMTS12, ADGRL3, AMIGO3, ASS1, ASTN2, CDH13, CDH17, CDH19, CDH20, CDH23, CHL1, CLDN23, CNTN4, COL8A1, CTNNA1, CTNNA3, DSCAML1, DTX1, DYSF, EGFLAM, EPHA8, GPC4, IGSF11, ITGA11, ITGB2, LAMA3, LSAMP, MACF1, MAD1L1, MEGF9, MYH9, MYOT, NCAM1, NEGR1, NFKBIZ, NRXN1, PARD3B, PARVA, PCDH15, PLXNA1, PLXNA2, PLXNA4, PPP3CA, RDX, RET, STAB2, STK10, TLN1, TMEFF2, TNC, VTN, VWF, ZDHHC2 |
| GO:0003073 | regulation of systemic arterial blood pr... | 39 | 6 | 2.68 | 0.003 | ADRA1D, ADRB2, AR, NCALD, REN, WNK1 |
| GO:0001967 | suckling behavior | 7 | 3 | 0.48 | 0.009 | APP, CNTFR, UBE2Q1 |
| GO:0001778 | plasma membrane repair | 7 | 3 | 0.48 | 0.009 | DYSF, MYH9, SYT11 |

**Table S10:**
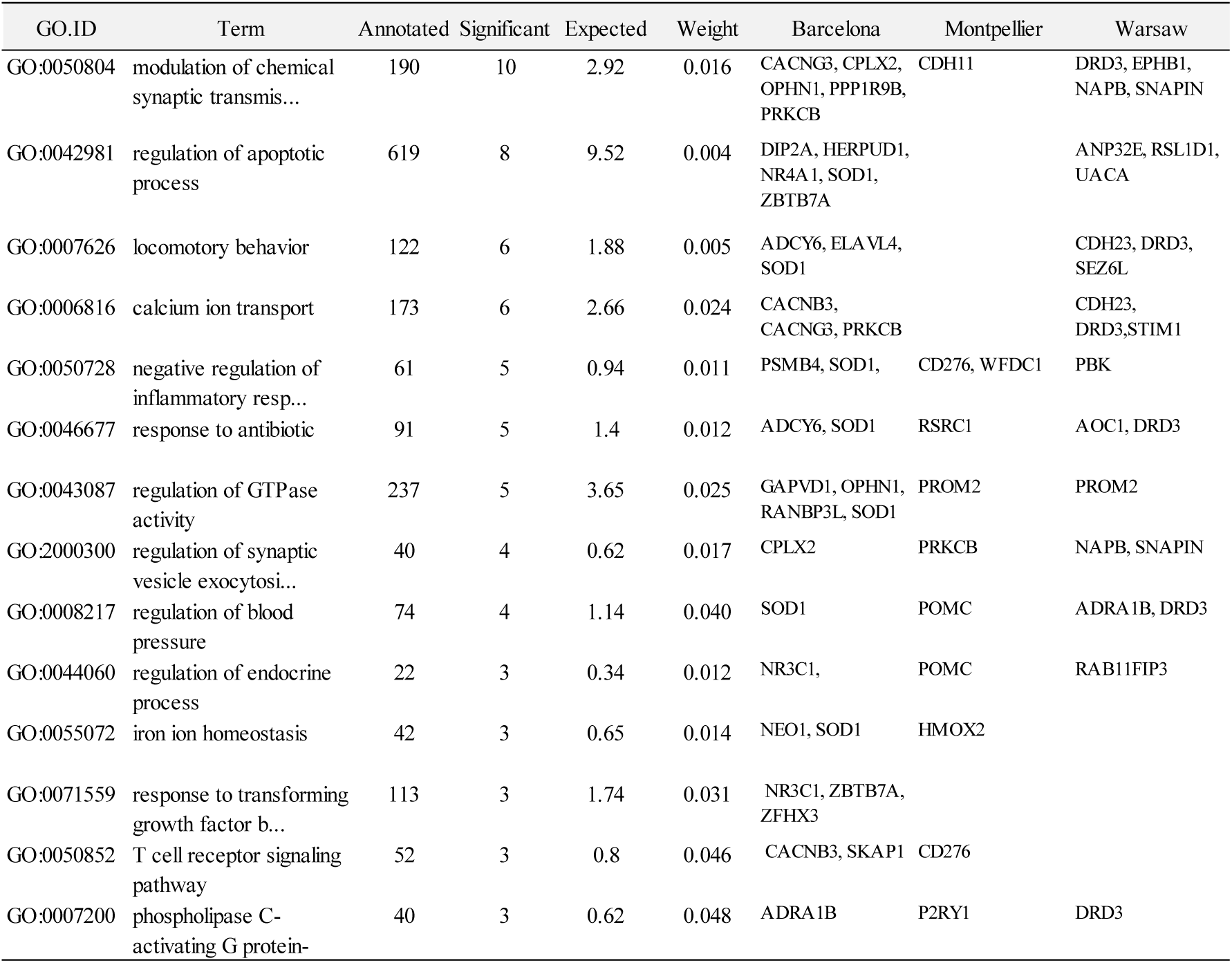
Significantly enriched GO associated with genes overlapping 5kb windows around DMRs between forest and urban habitats.

**Table S11:**
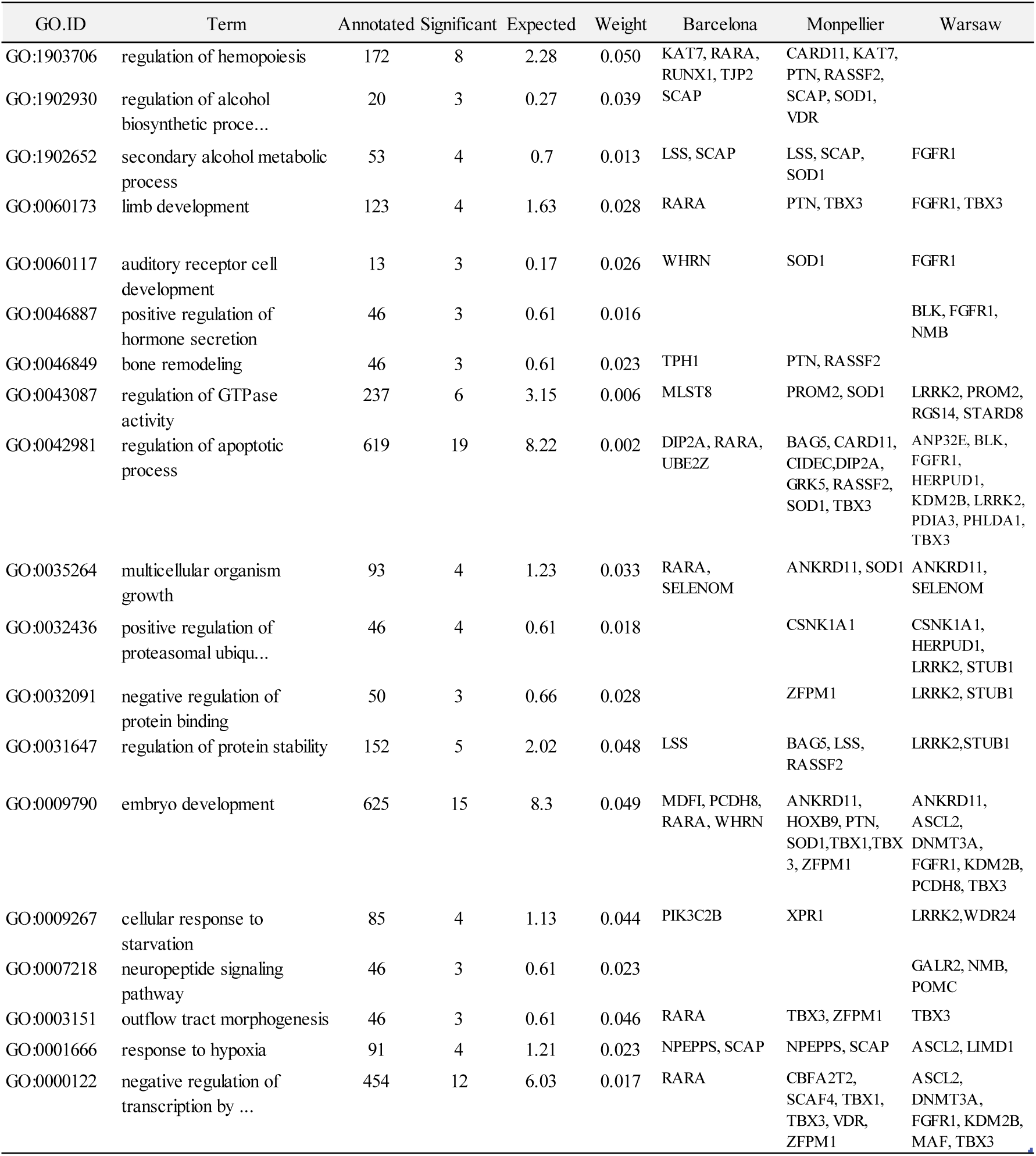
Significantly enriched GO associated with genes overlapping 5kb windows around DMRs between sexes.

**Table S12:** Number of DMRs between habitats, found in females and males separately. See Supplementary Analysis 2. “Hypermethylated” and “Hypomethylated” refer to hyper- or hypomethylated DMR in the urban habitat compared to the forest habitat.

|  | Females |  |  |
| --- | --- | --- | --- |
|  | Barcelona | Montpellier | Warsaw |
| Hypermethylated | 240 | 275 | 205 |
| Hypomethylated | 224 | 133 | 316 |
| Total | 464 | 408 | 521 |
|  | Males |  |  |
|  | Barcelona | Montpellier | Warsaw |
| Hypermethylated | 107 | 347 | 315 |
| Hypomethylated | 317 | 229 | 218 |
| Total | 424 | 576 | 533 |

**Table S13:**
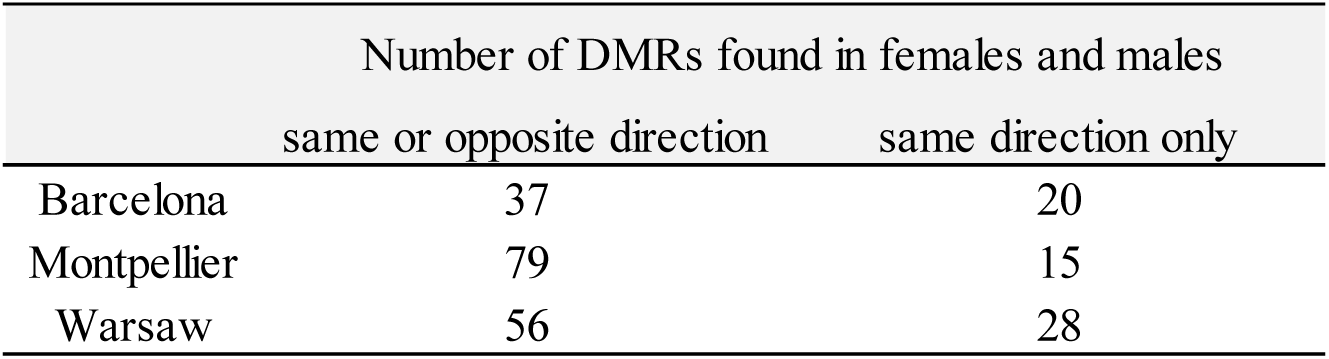
Number of habitat linked DMRs shared between females and males. See Supplementary Analysis 2. “Same direction” refers to hyper- or hypo-methylated in both sexes.

|  | Number of DMRs found in females and males |  |
| --- | --- | --- |
|  | same or opposite direction | same direction only |
| Barcelona | 37 | 20 |
| Montpellier | 79 | 15 |
| Warsaw | 56 | 28 |

### SUPPLEMENTARY FIGURES

**Figure S1:**
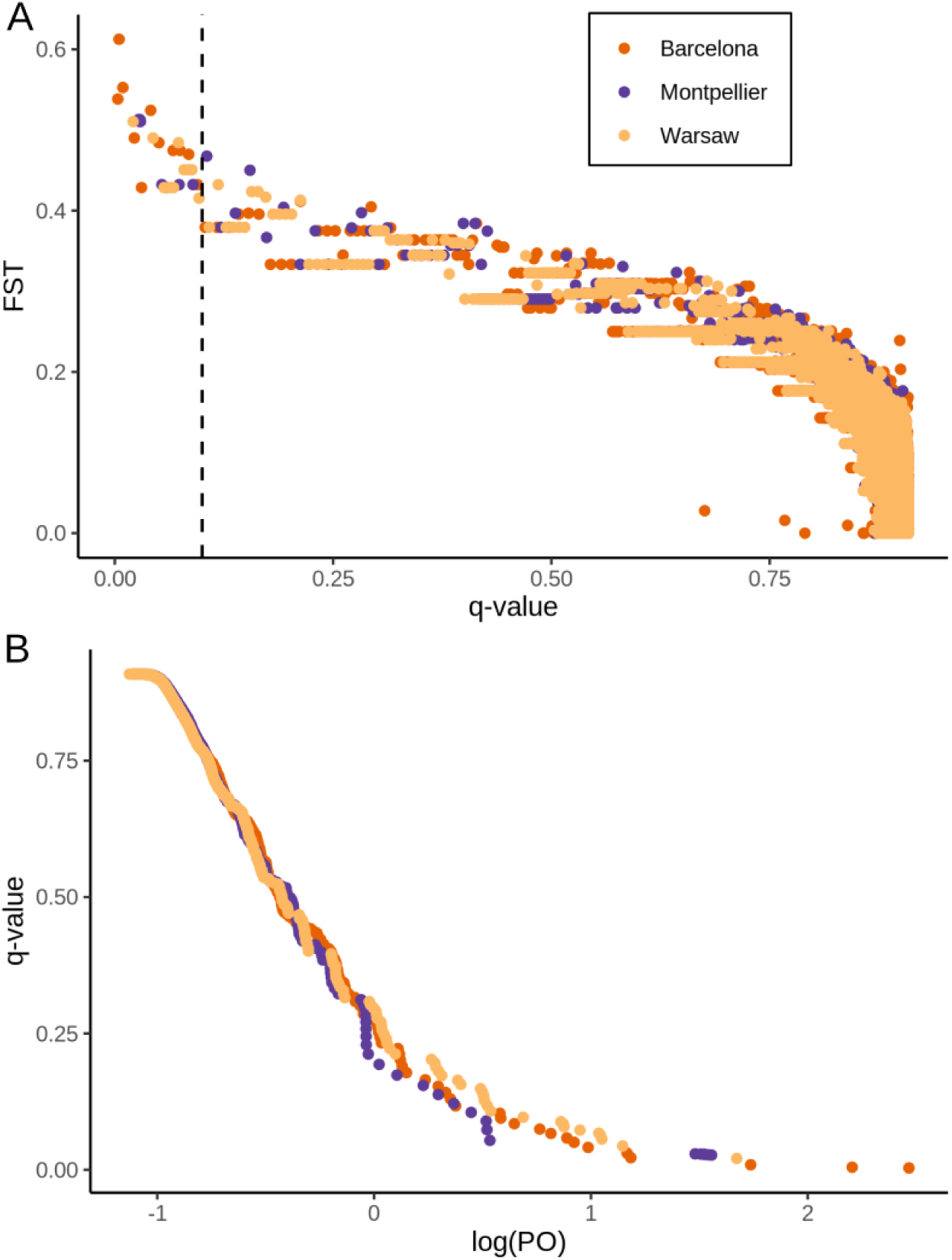
Results of the Bayescan tests between forest and urban great tits, for each of the three urban-forest pairs. (A) Distribution of F_ST_ values per position in function of q-value. (B) Distribution of q-value in function of log(PO).

**Figure S2:**
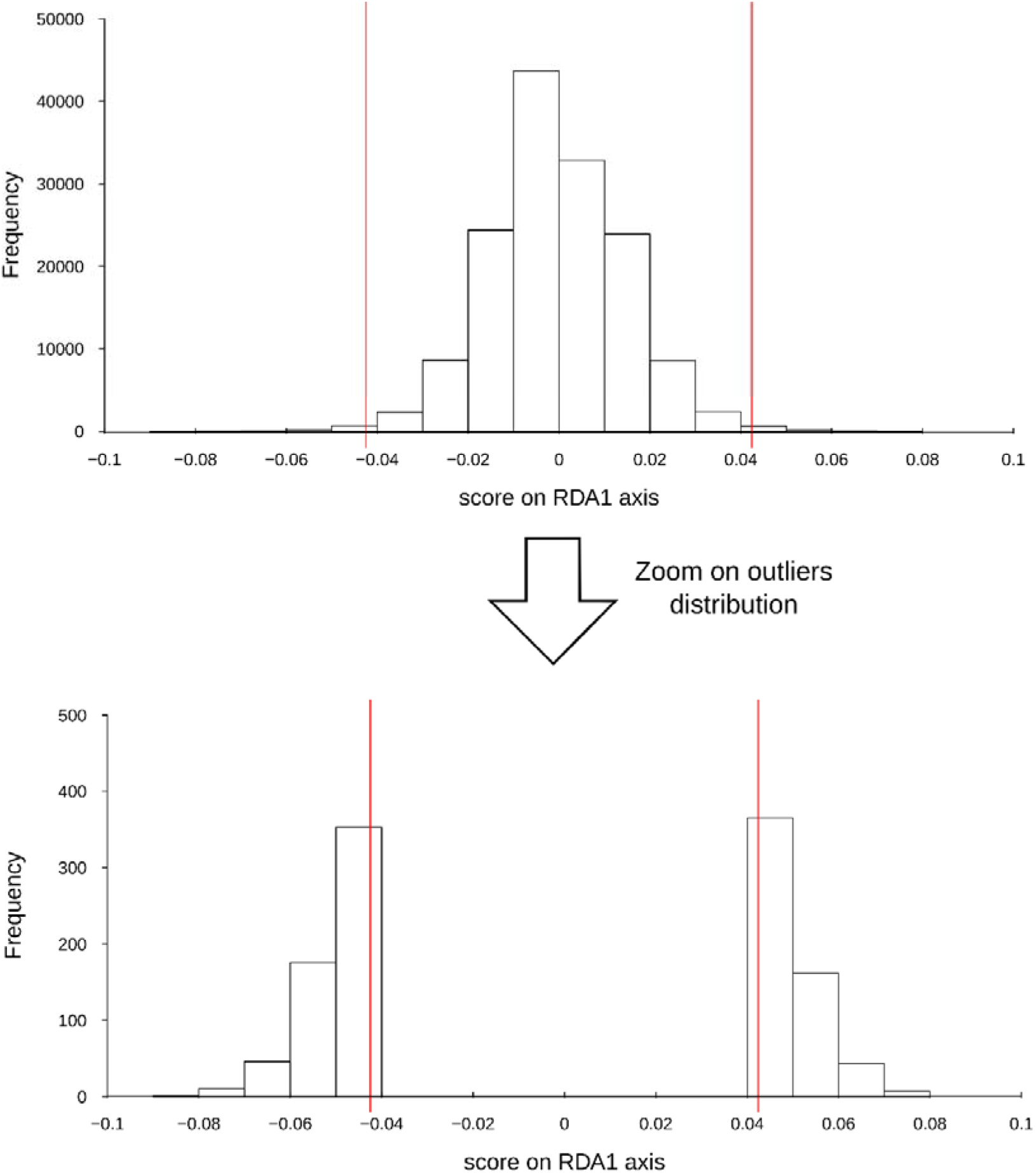
Distribution of genomic RDA loadings of each locus (above), and outliers (below). Vertical red lines represent the thresholds used to select outliers (please see methods for more details).

**Figure S3:**
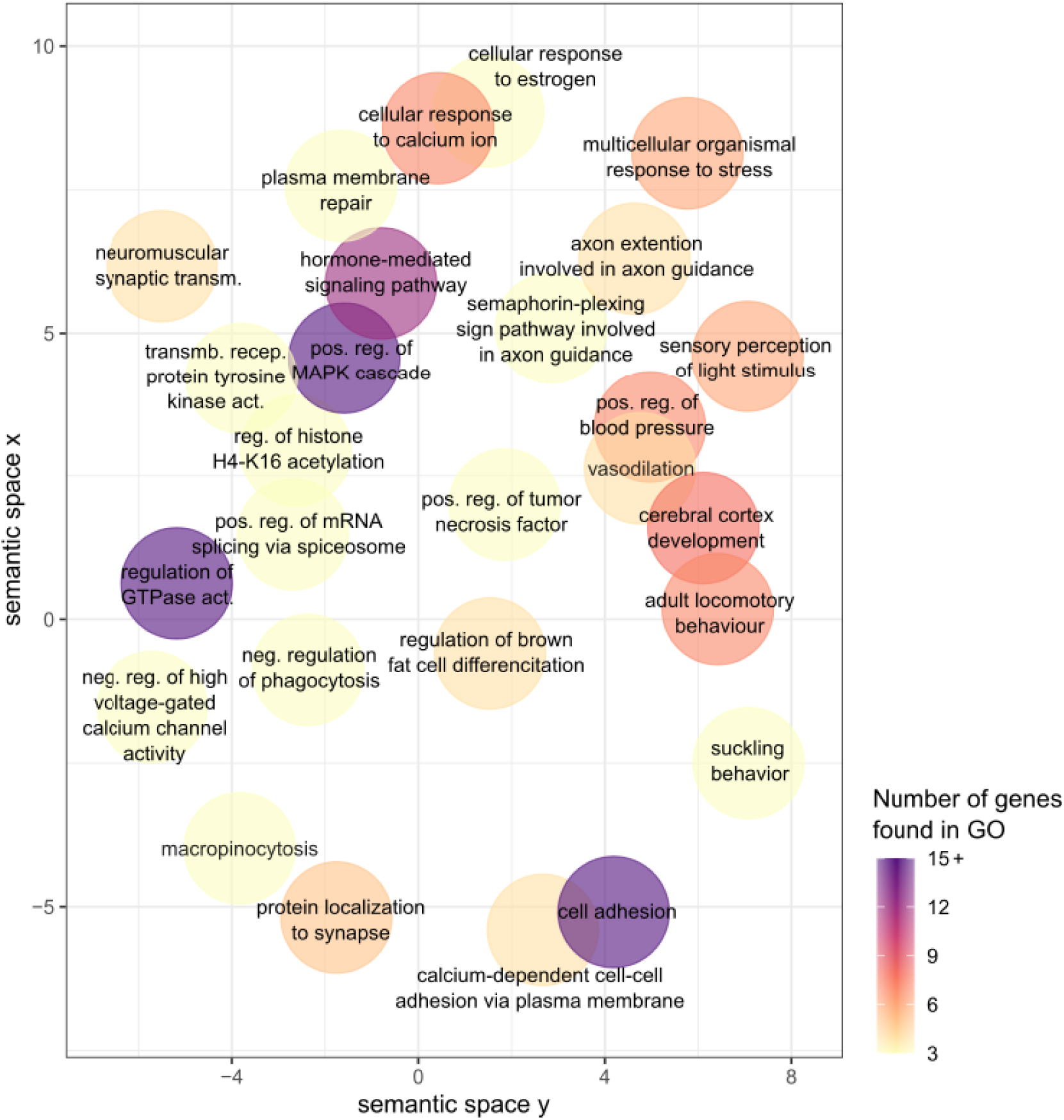
Semantic space representation for enriched GO terms from genes associated with genetic outlier (the outliers of the RDA analyses). Semantic space coordinates have been calculated based on similarity of GO term word composition using REVIGO. The color gradient indicates the number of genes involved in each GO term.

**Figure S4:**
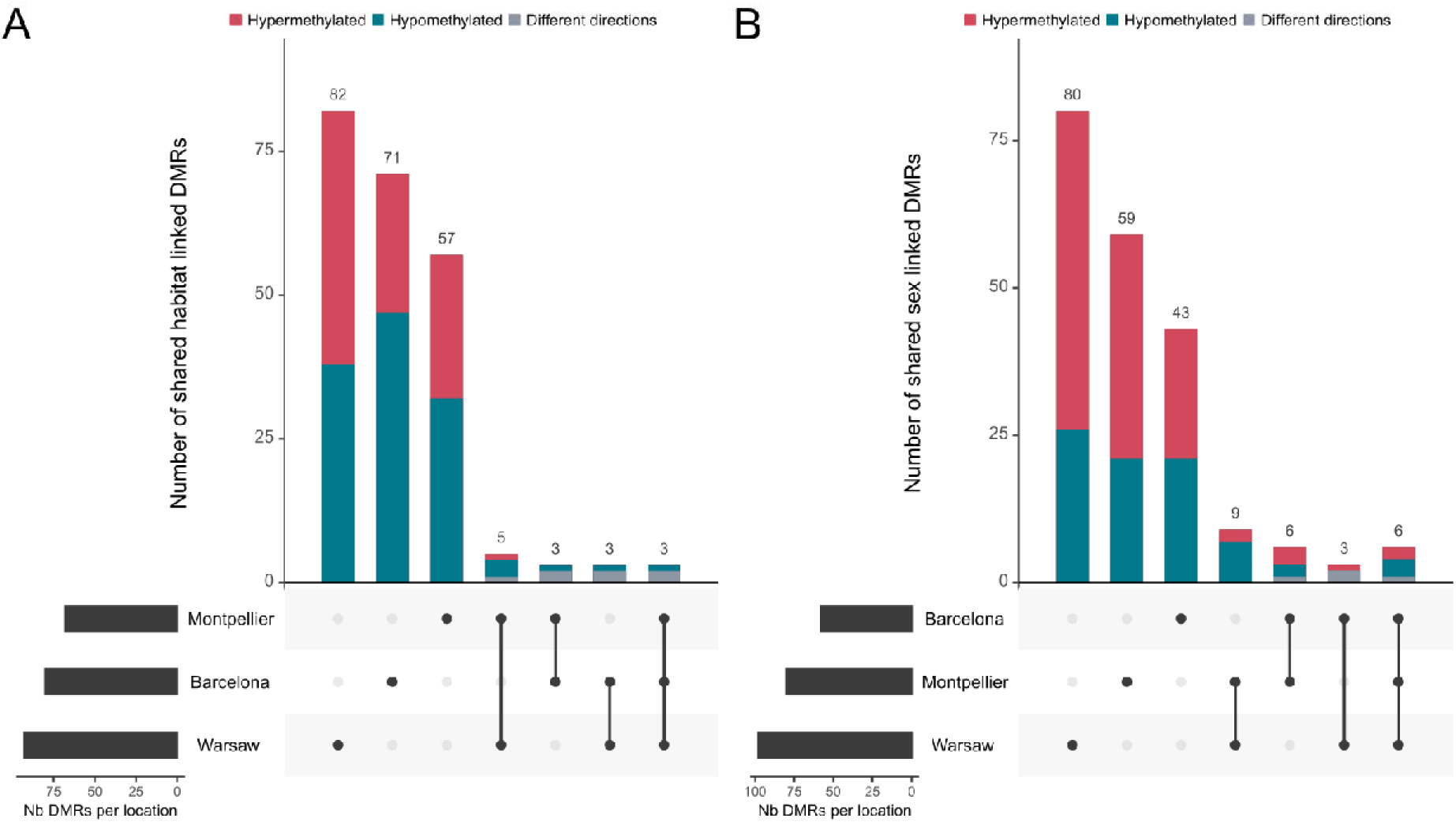
Sharing among each location of the significant DMRs found between A) forest and urban habitats, and B) females and males great tits. Hypermethylated DMRs (A) in urban, B) in males) are shown in red, hypomethylated in blue and for shared DMRs, grey represent cases where a DMRs was found in multiple locations but for which the direction of methylation was opposite.

**Figure S5:**
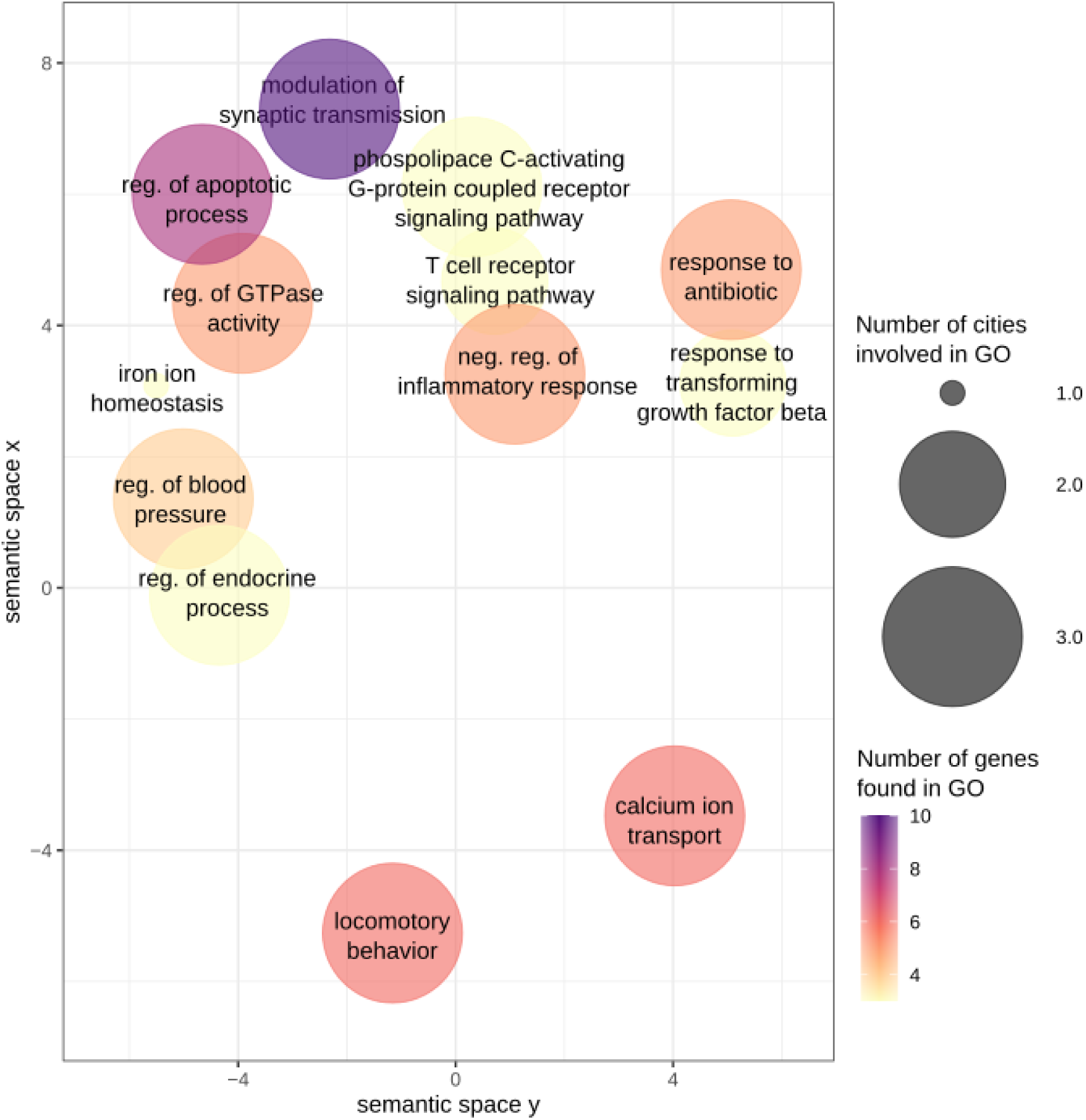
Semantic space of enriched GO terms from genes associated with urbanization-linked DMRs. Semantic space coordinates have been calculated based on similarity of GO term word composition using REVIGO. The color gradient indicates the number of genes involved in each GO term and circle size indicates the number of city contributing to each category.

**Figure S6:**
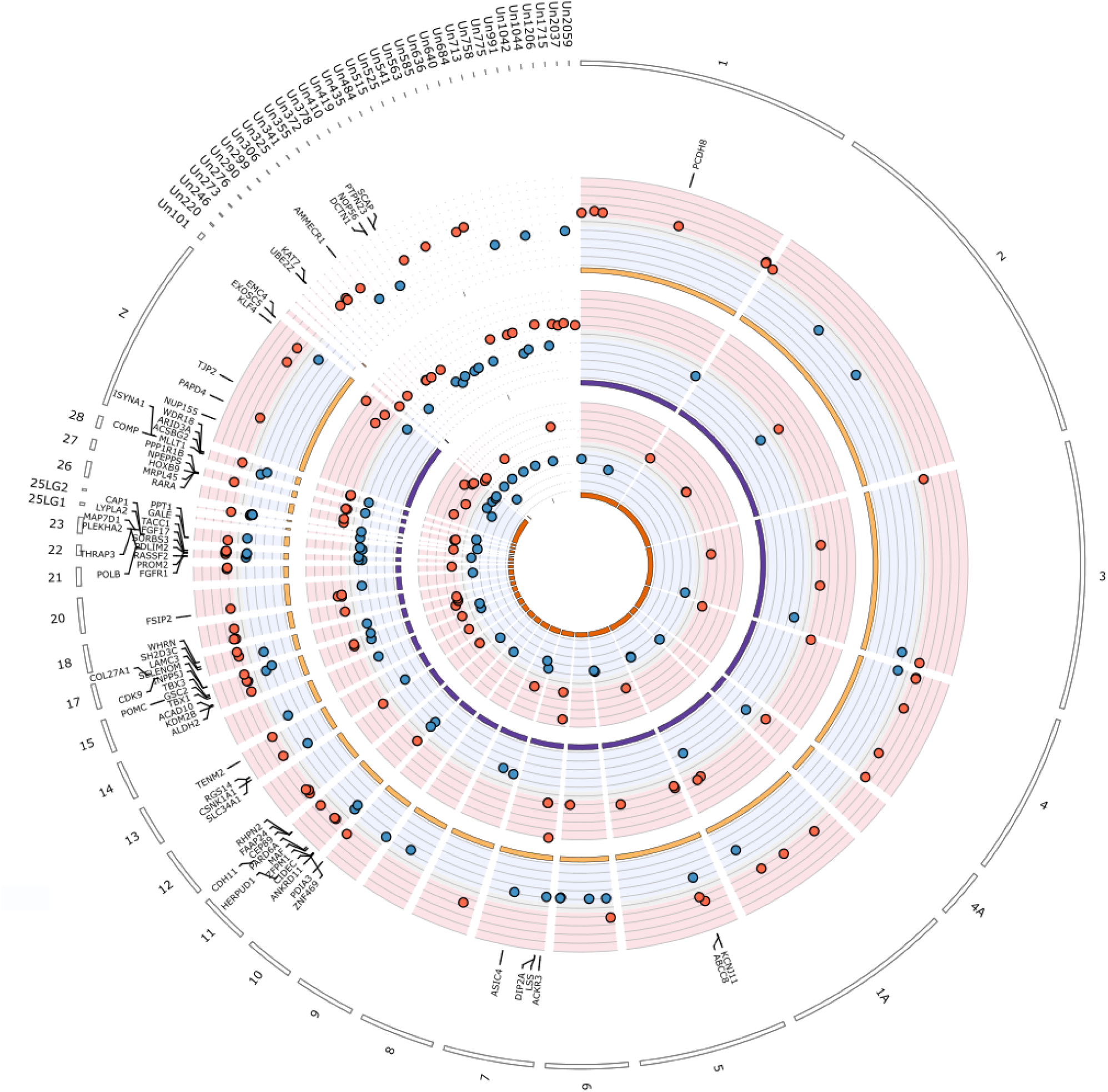
Circos plot of differentially methylated regions (DMRs) identified between females and males great tits in and near Barcelona, Montpellier and Warsaw (from inner to outer circles). Red points show hypermethylated regions in female great tits relatively to males, and blue points show hypomethylated regions. For graphical clarity, only a subset of genes are represented: genes associated with the 10% most extreme DMR. Names of the genes found within 5 kb of the represented DMRs are given.

**Figure S7:**
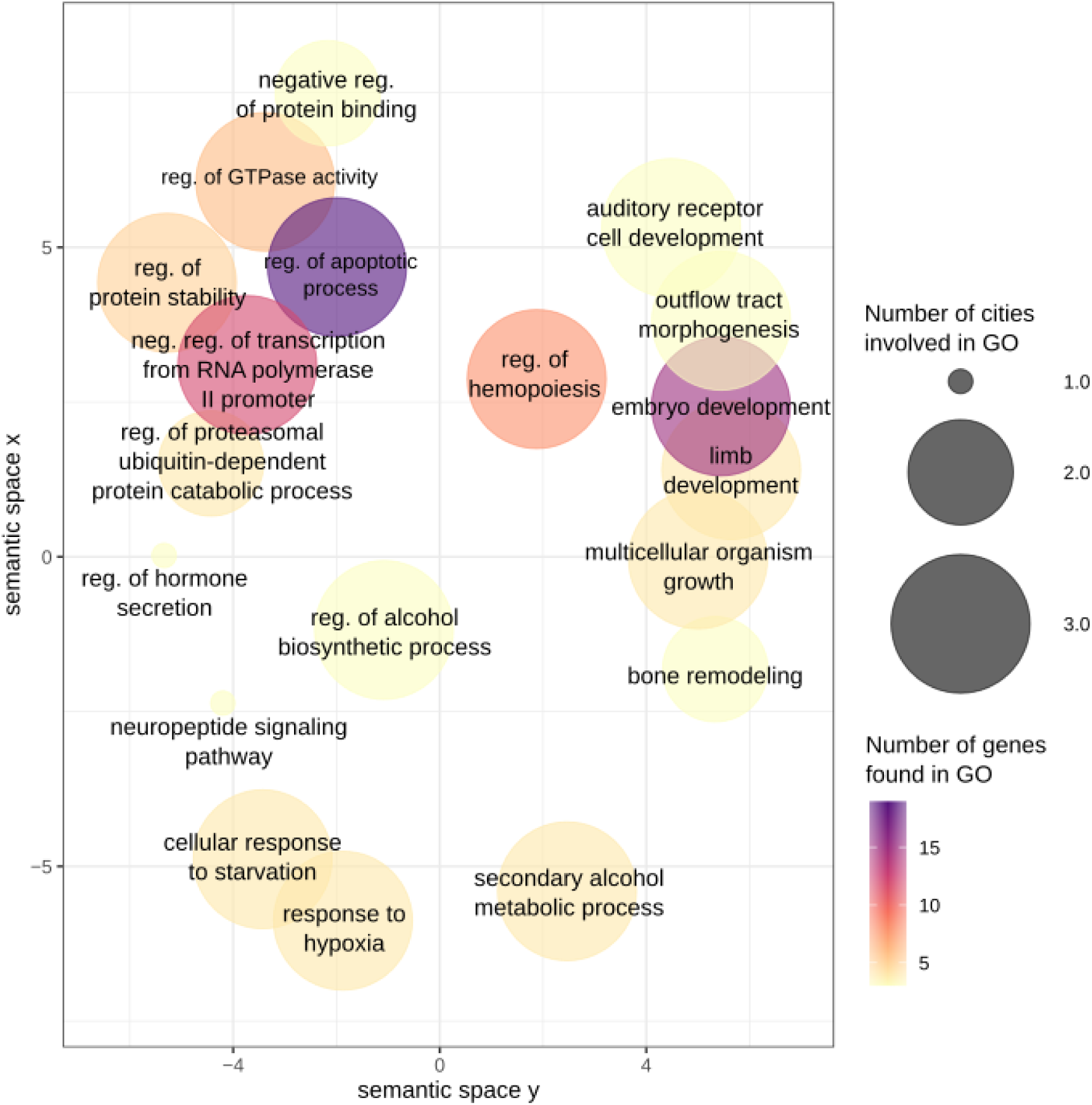
Semantic space of enriched GO terms from genes associated with sex-linked DMRs. Semantic space coordinates are calculated based on similarity of GO term word composition using REVIGO. The color gradient indicates the number of genes involved in each GO term and circle size indicates the number of city contributing to each category.

**Figure S8:**
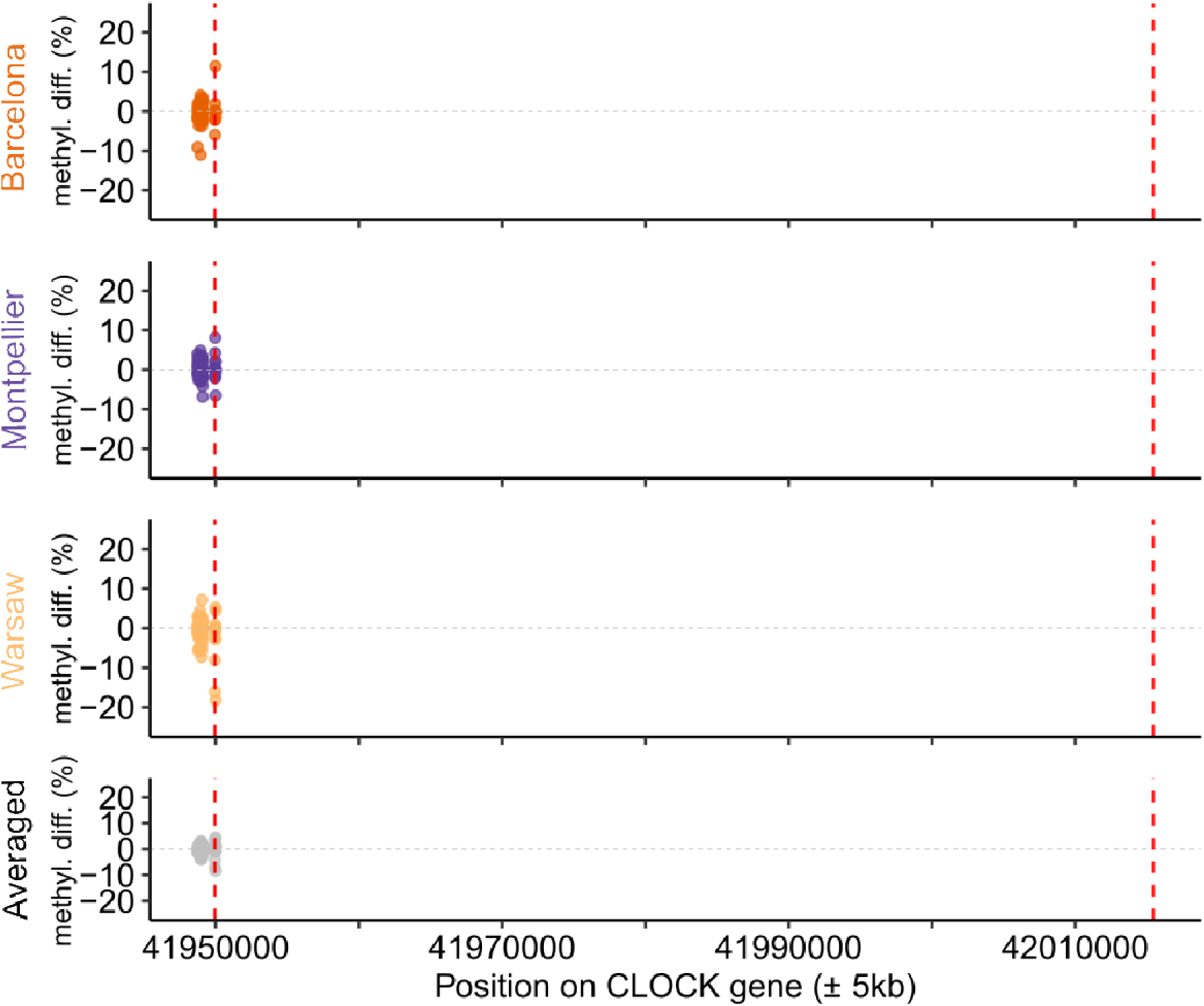
Mean methylation difference per base along the CLOCK gene. Vertical dashed lines materialised start and end of the gene.

**Figure S9:**
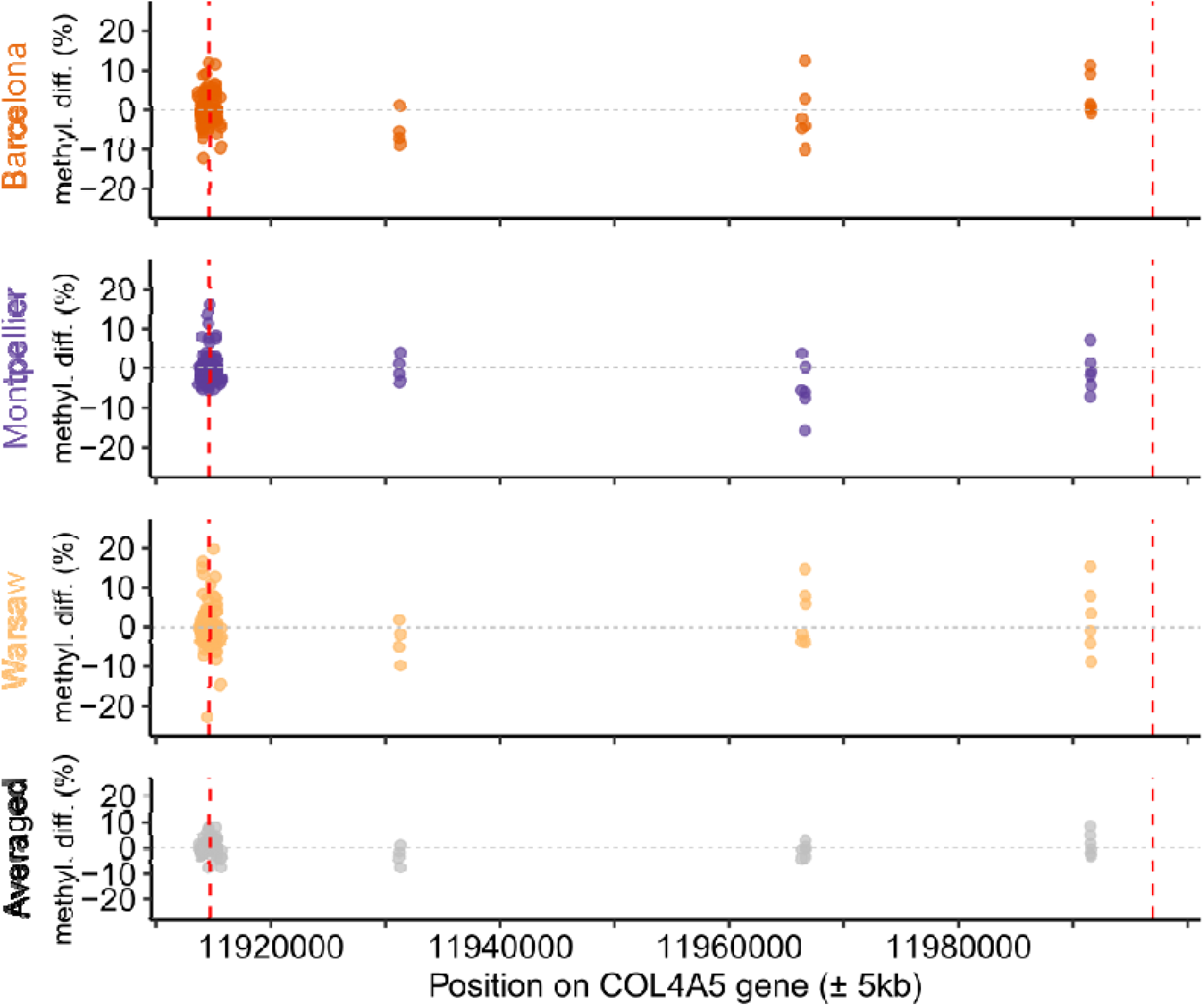
Mean methylation difference per base along the COL4A5 gene. Vertical dashed lines materialised start and end of the gene.

**Figure S10:**
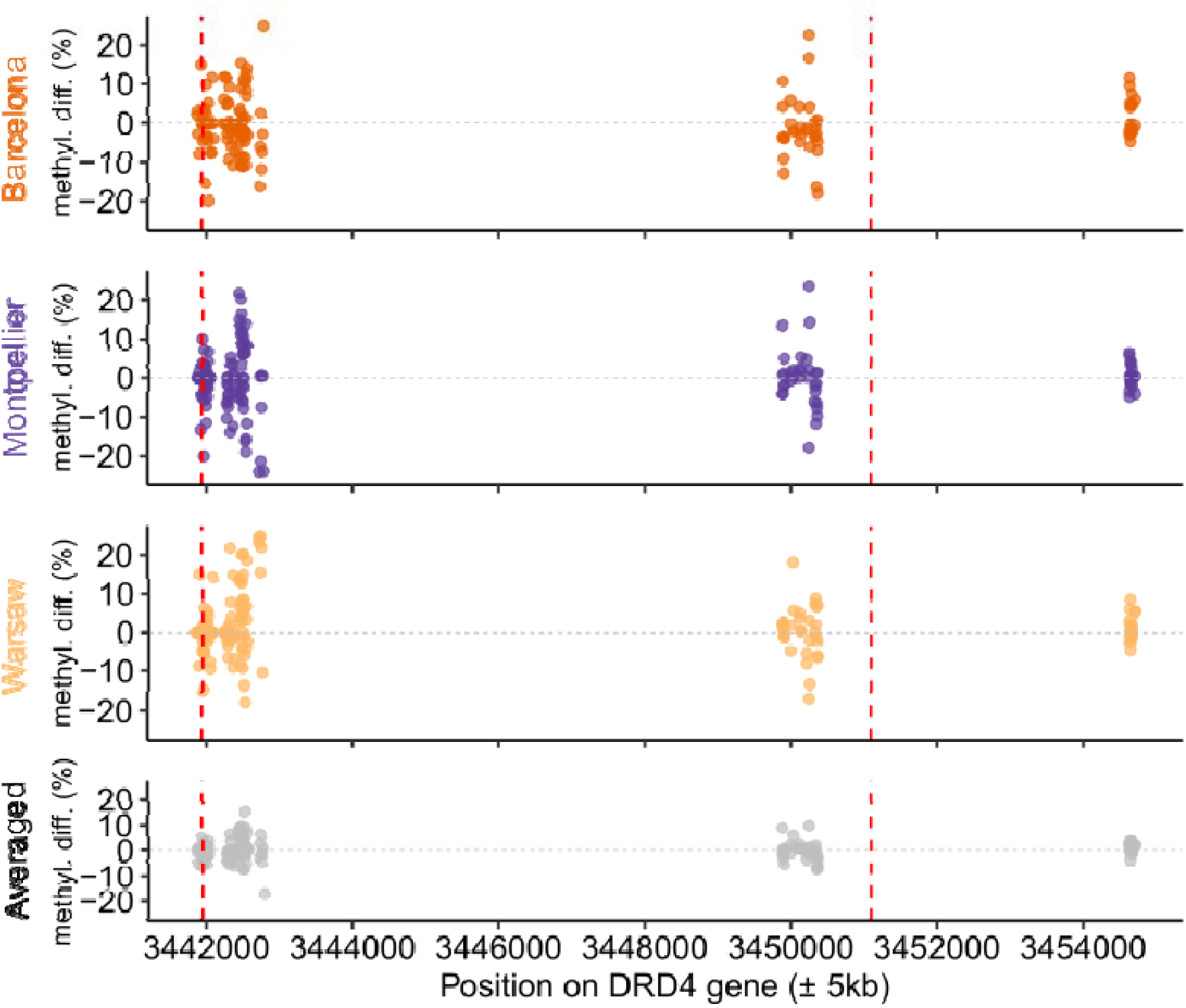
Mean methylation difference per base along the DRD4 gene. Vertical dashed lines materialised start and end of the gene.

**Figure S11:**
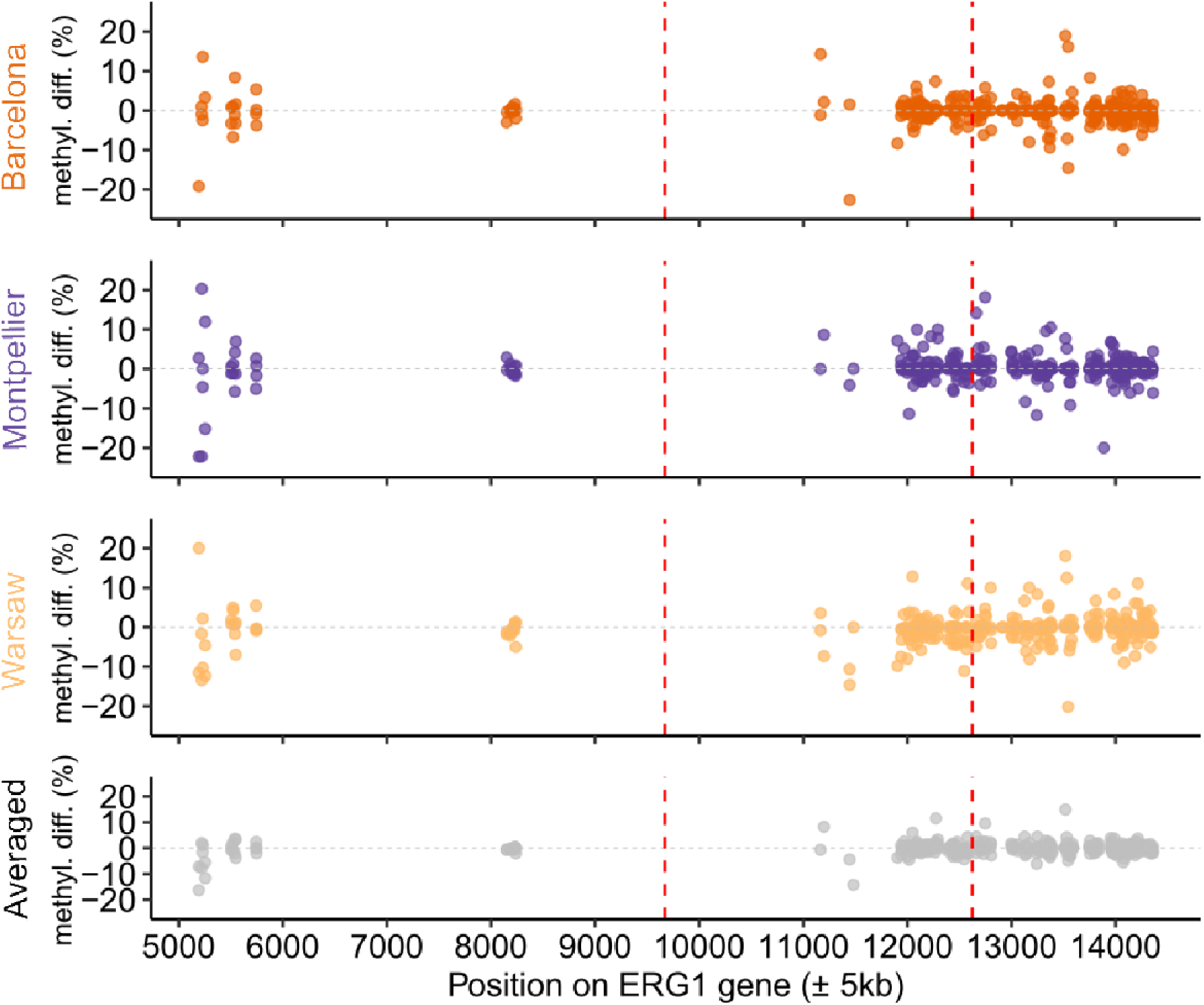
Mean methylation difference per base along the ERG1 gene. Vertical dashed lines materialised start and end of the gene.

**Figure S12:**
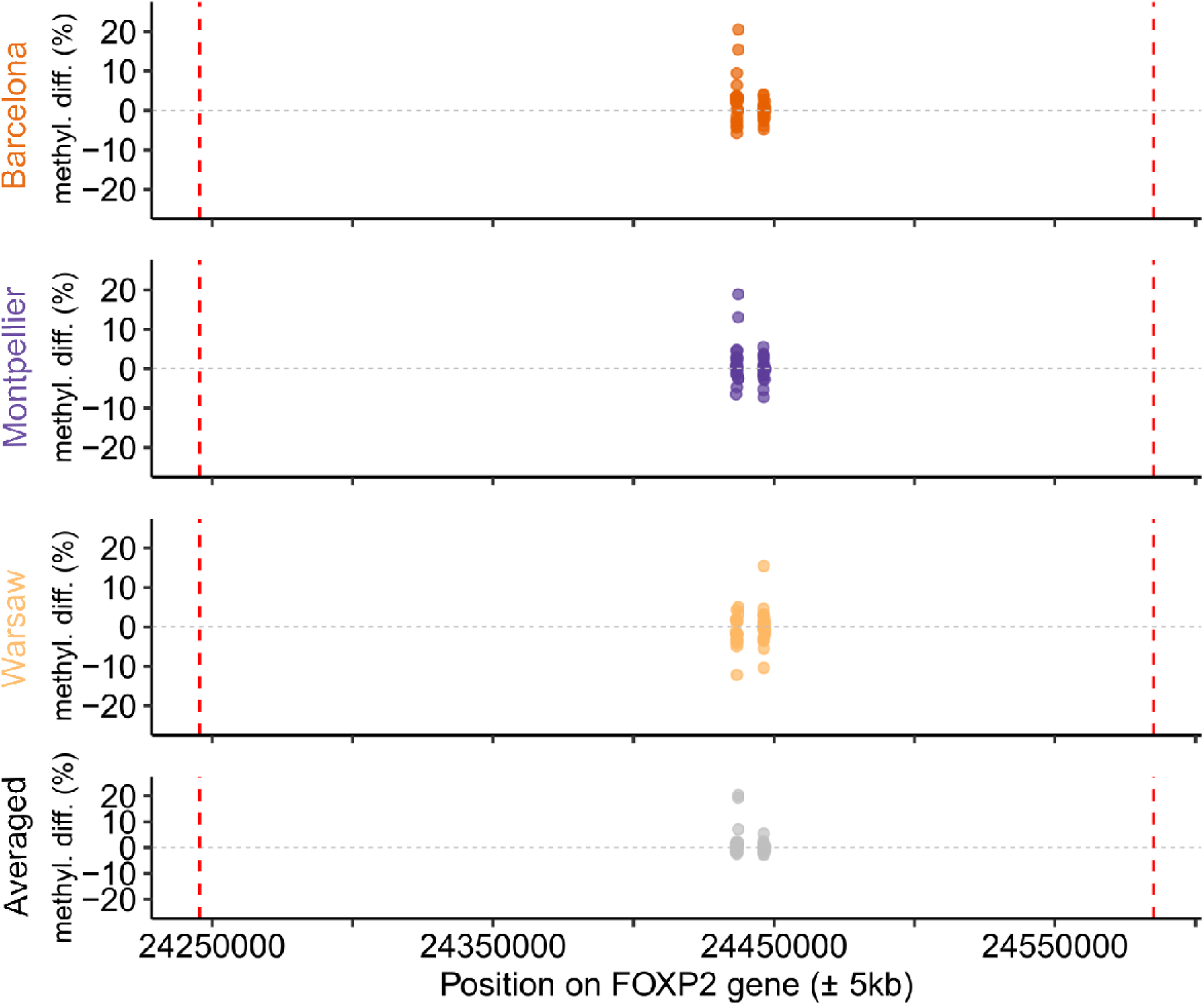
Mean methylation difference per base along the FOXP2 gene. Vertical dashed lines materialised start and end of the gene.

## SUPPLEMENTARY ANALYSES

### Differentially Methylated Regions (DMRs) between Females and Males

A total of 238 DMR were found between sexes: 58 in Barcelona, 81 in Montpellier and 99 in Warsaw (Figure S6). Warsaw presented significantly more hyper than hypomethylated DMR (χ^2^=5.878, P= 0.015), but it was not the case for Barcelona (χ^2^=0, P=1) nor Montpellier (χ^2^=1.25, P=0.264). DMR were distributed on 29 chromosomes and 35 unplaced scaffolds.

In total, 181 DMR were located on genes or in 5kb upstream/downstream regions around genes. A gene ontology enrichment analysis performed on the pooled list of genes associated with DMR revealed significantly enriched modules associated with metabolism, development and hormonal function (see details Figure S4B and S6, and Table S11).

### Differentially Methylated Regions between associated with urbanization in females and males

To explore the potential sex dependent effects of urbanization on methylation patterns, we searched for DMRs between forest and urban habitats in females and males separately. The numbers of significant DMRs per sex per location are reported in Table S12.

In females, 101 DMRs (7.9%) were shared between 2 or more locations and 45 (3.5%) showed the same direction of variation in all cities.

Similarly, in males 134 DMRs (9.7%) were shared by multiple locations, and 58 (4.18%) showed the same direction of variation.

Within locations, sexes showed shared urban-linked DMRs, see Table S12.

