## Supplementary information for "Testing for parallel genomic and epigenomic footprints of adaptation to urban life in a passerine bird"

**SUPPLEMENTARY TABLES**

**Table S1:** Redundancy analysis (RDA) performed on the genetic data including the Z chromosome.

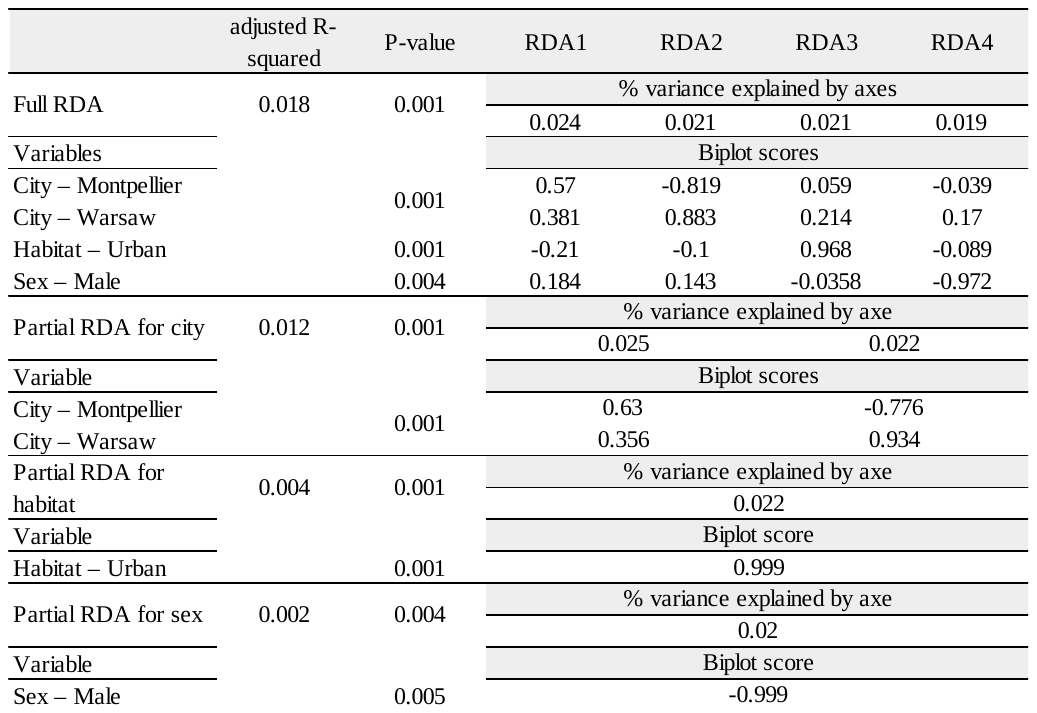

**Table S2:** Redundancy analysis (RDA) performed on the genetic data without Z chromosome.

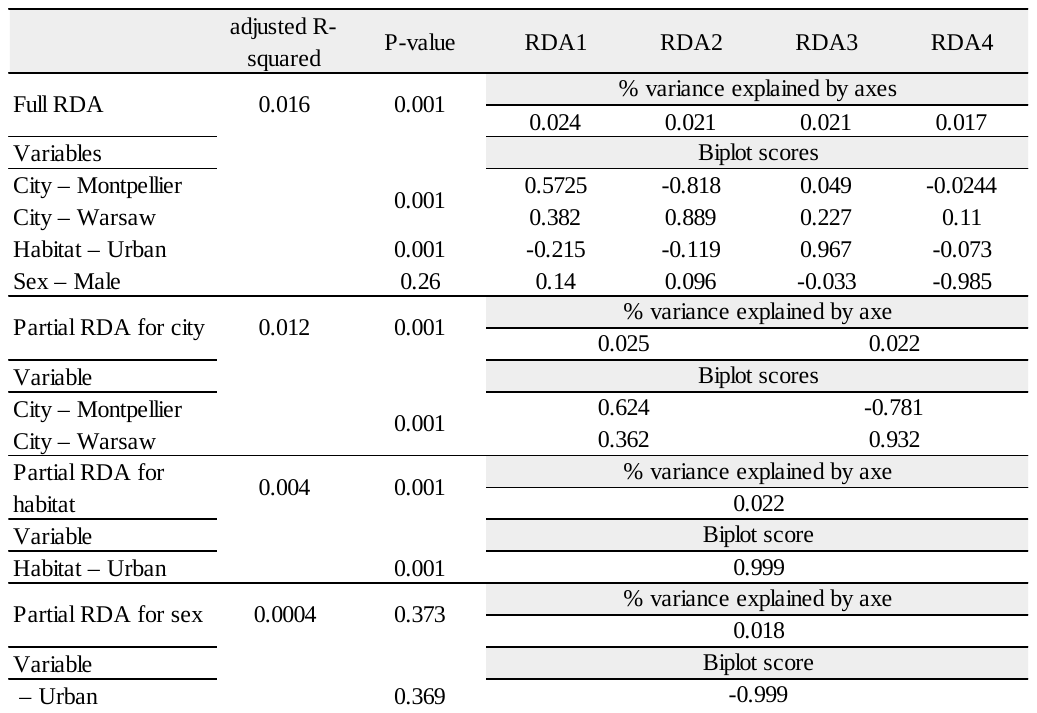

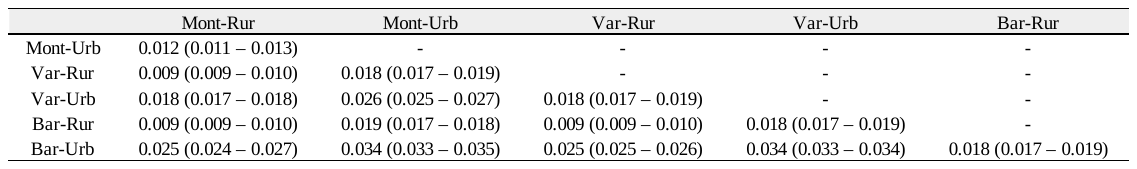
**Table S3:** Fst estimation between pairs of subpopulations. 95% confidence computed intervals in brackets were computed using StAMPP package with 1000 bootstrap.

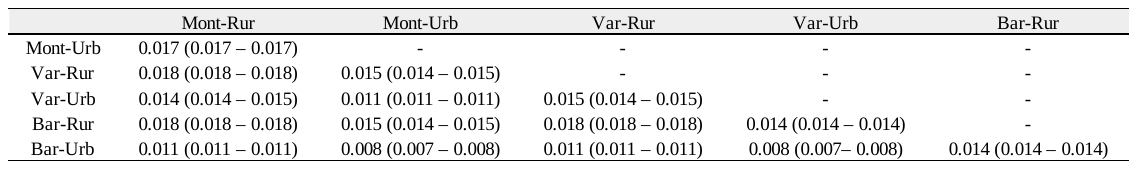
**Table S4:** Fst estimation averaged on autosomes between pairs of subpopulations. 95% confidence computed intervals in brackets were computed using StAMPP package with 1000 bootstrap.

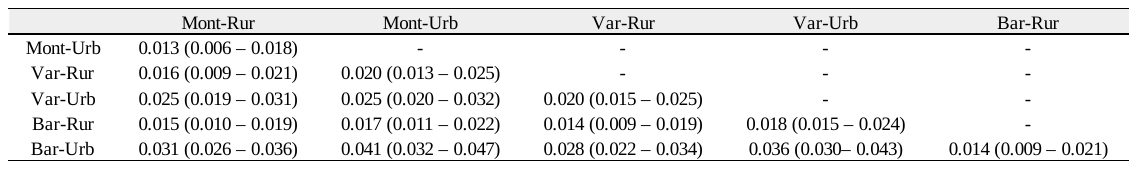
**Table S5:** Fst estimation averaged on Z chromosome between pairs of subpopulations. 95% confidence computed intervals in parenthesis were computed using StAMPP package with 1000 bootstrap.

**Table S6:**  Redundancy analysis (RDA) performed on the methylation data including the Z chromosome.

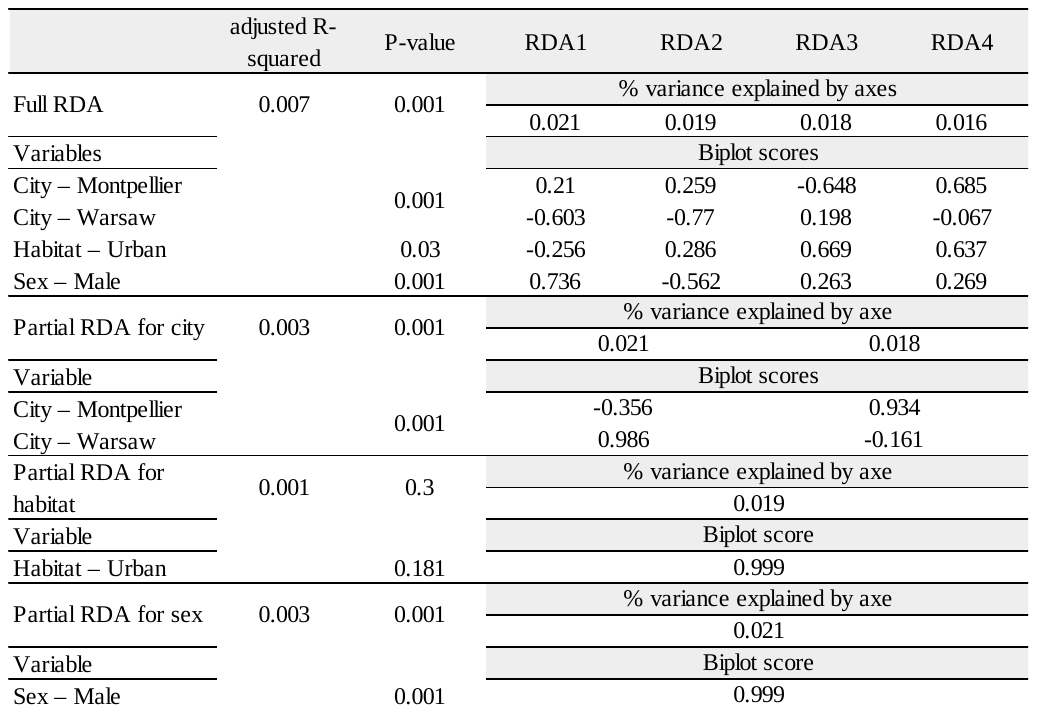

**Table S7:**  Redundancy analysis (RDA) performed on the methylation data without Z chromosome.

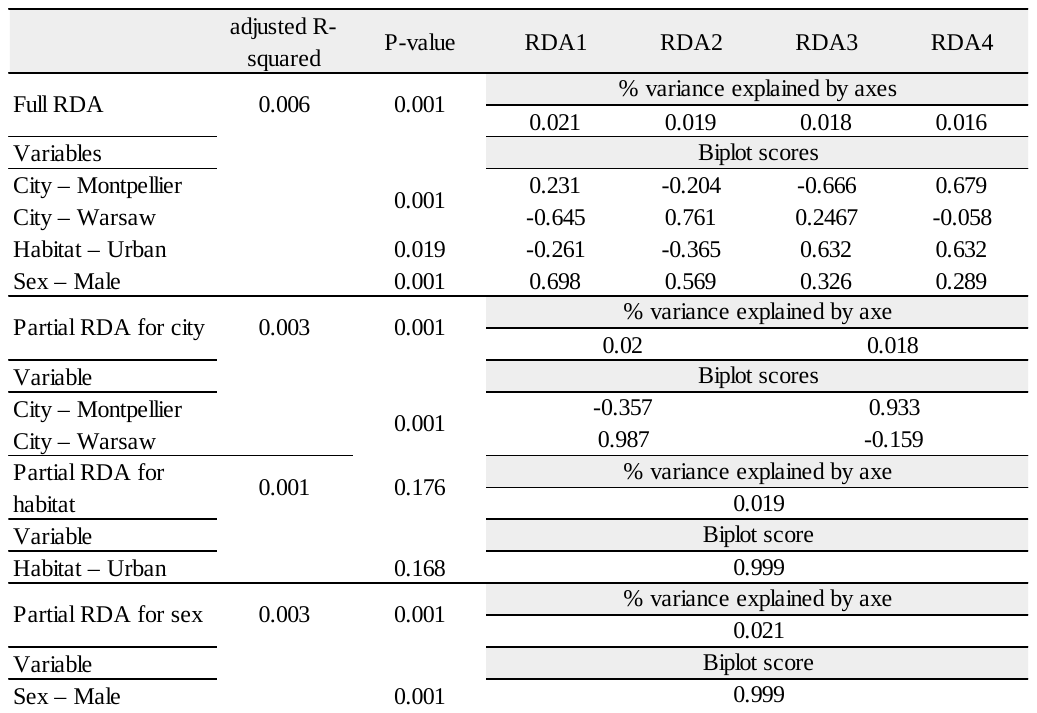

**Table S8:** Results of Type I ANOVA performed on mean methylation level of cytosines in a CpG context per individual, including: habitat (rural vs urban), city (Barcelona, Montpellier and Warsaw) and sex as explanatory variables. The first part of the table shows results of the analysis on autosomes and the second part shows results of the analysis on the Z chromosome only.

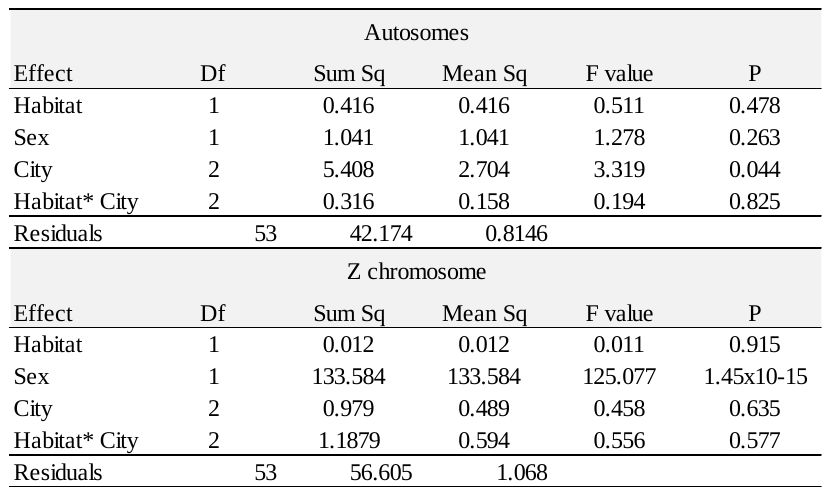

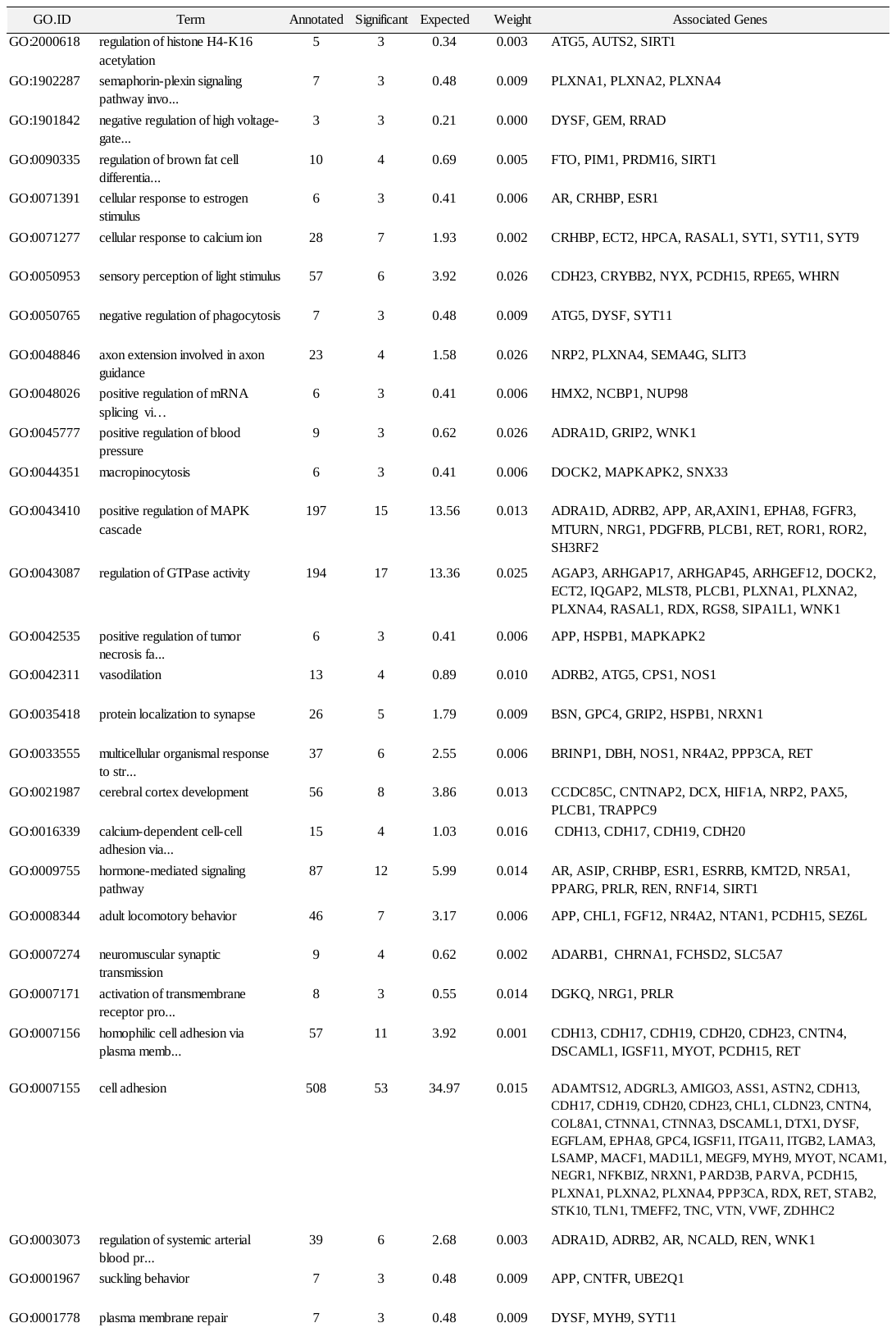
**Table S9:** Significantly enriched GO associated with genes overlapping 5kb windows around genomic outliers between forest and urban habitats.

**Table S10:** Significantly enriched GO associated with genes overlapping 5kb windows around DMRs between forest and urban habitats.

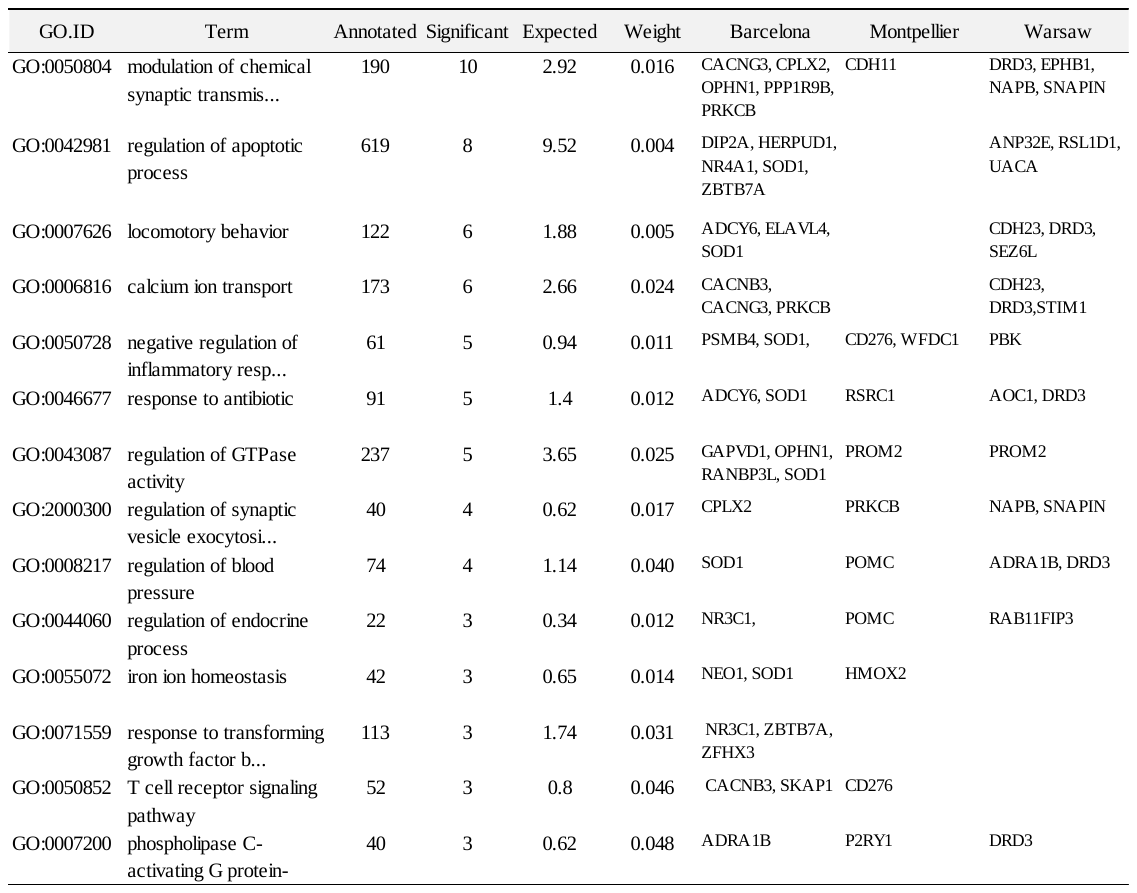

**Table S11:** Significantly enriched GO associated with genes overlapping 5kb windows around DMRs between sexes.

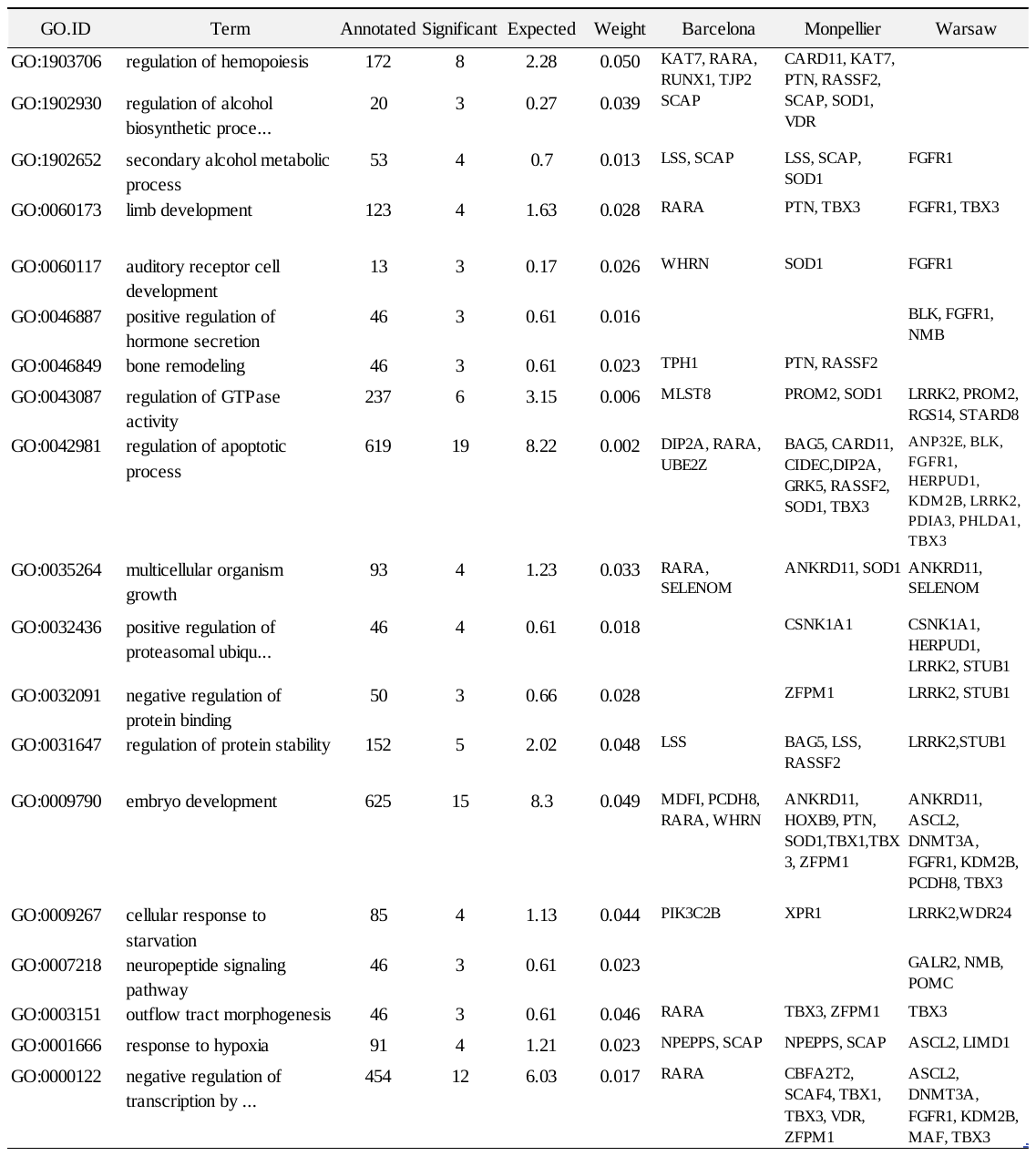

**Table S12:** Number of DMRs between habitats, found in females and males separately. See Supplementary Analysis 2. “Hypermethylated” and “Hypomethylated” refer to hyper- or hypomethylated DMR in the urban habitat compared to the forest habitat.

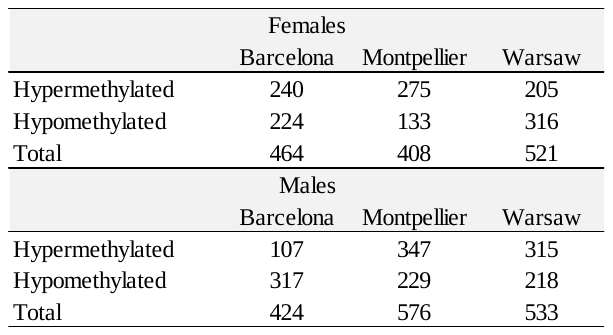

**Table S13:** Number of habitat linked DMRs shared between females and males. See Supplementary Analysis 2. “Same direction” refers to hyper- or hypo-methylated in both sexes.

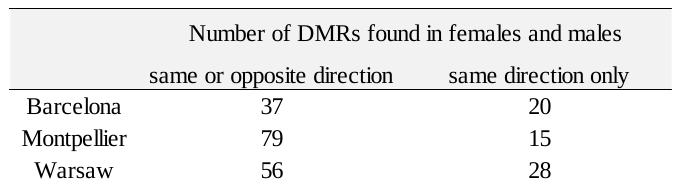

**SUPPLEMENTARY FIGURES**

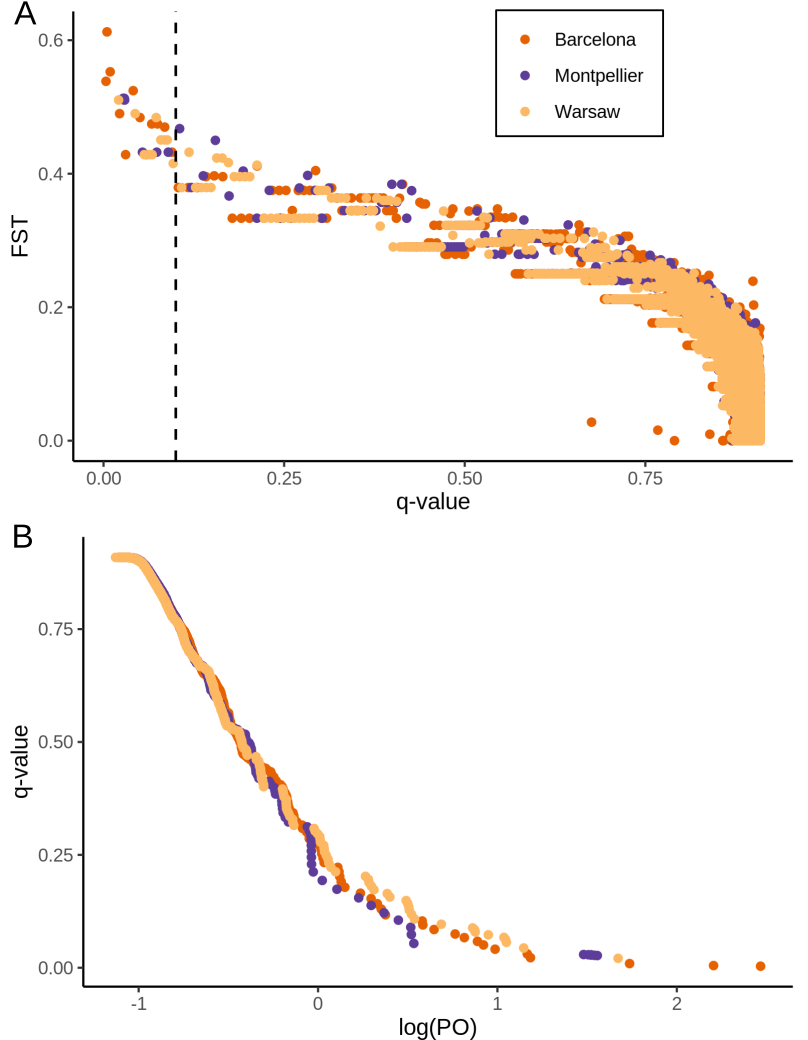

**Figure S1:** Results of the Bayescan tests between forest and urban great tits, for each of the three urban-forest pairs. (A) Distribution of F_ST_ values per position in function of q-value. (B) Distribution of q-value in function of log(PO).

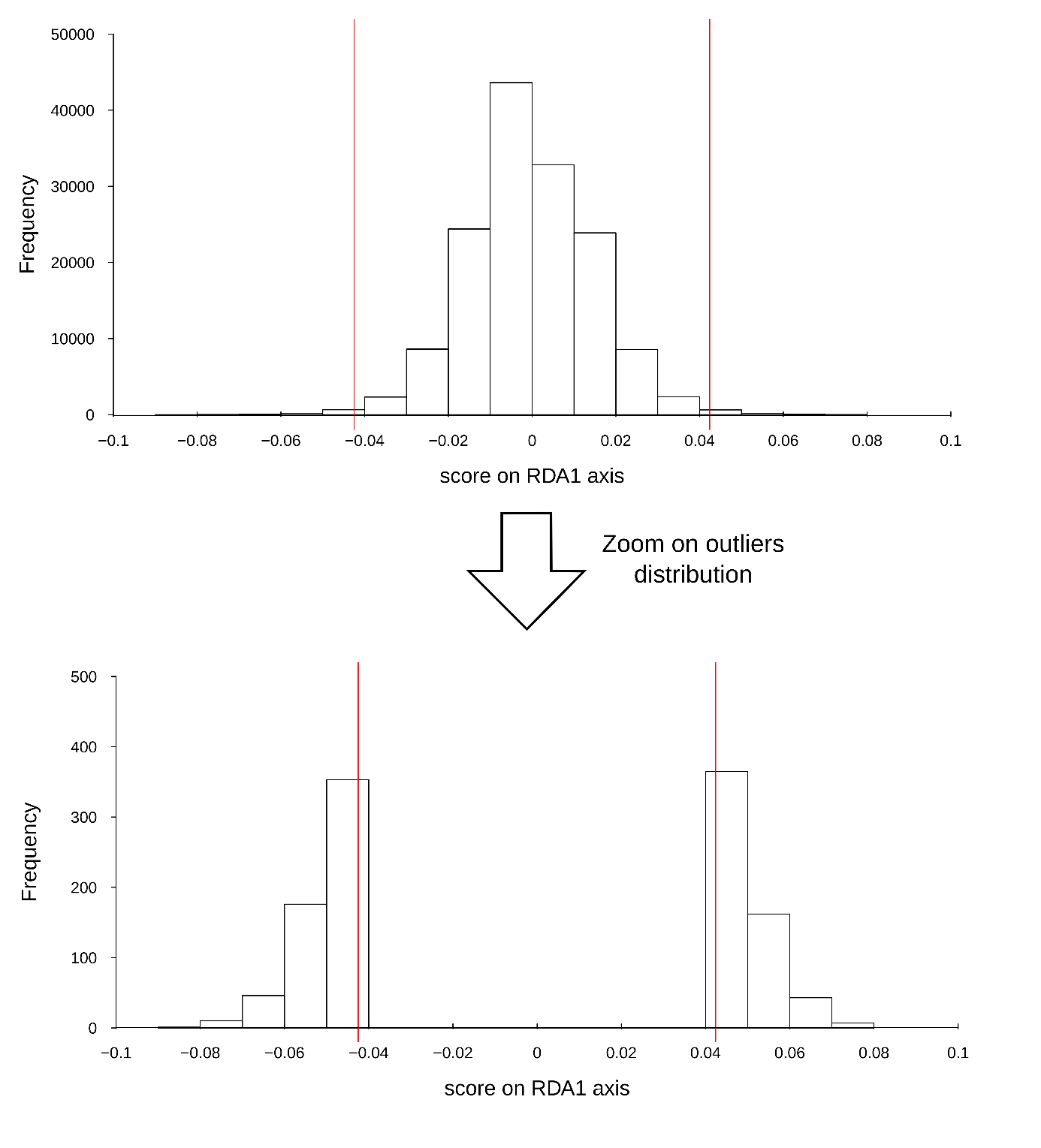

**Figure S2:** Distribution of genomic RDA loadings of each locus (above), and outliers (below). Vertical red lines represent the thresholds used to select outliers (please see methods for more details).

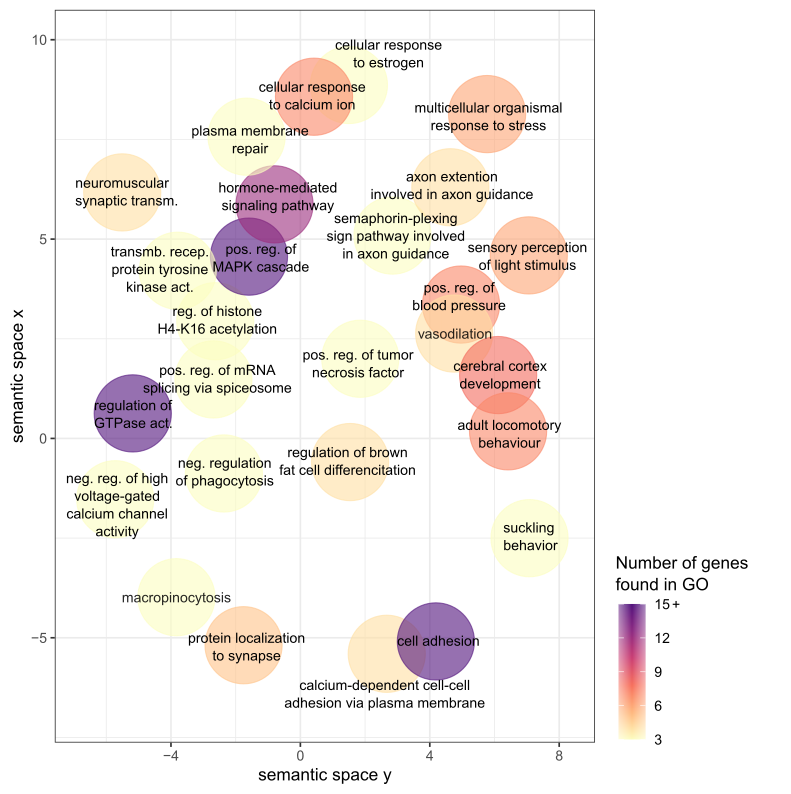

**Figure S3:** Semantic space representation for enriched GO terms from genes associated with genetic outlier (the outliers of the RDA analyses). Semantic space coordinates have been calculated based on similarity of GO term word composition using REVIGO. The color gradient indicates the number of genes involved in each GO term.

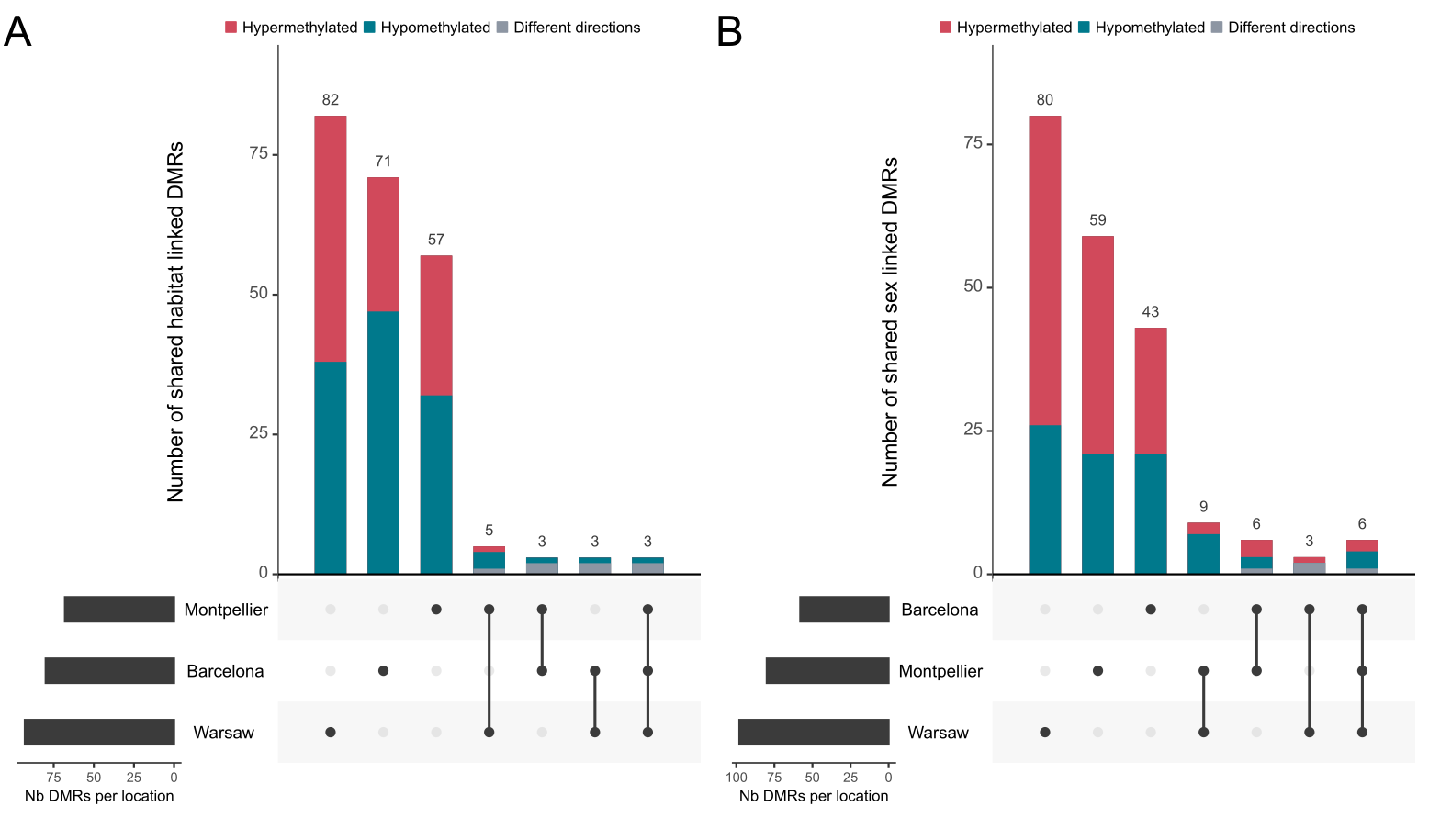
**Figure S4:** Sharing among each location of the significant DMRs found between A) forest and urban habitats, and B) females and males great tits. Hypermethylated DMRs ( A) in urban, B) in males) are shown in red, hypomethylated in blue and for shared DMRs, grey represent cases where a DMRs was found in multiple locations but for which the direction of methylation was opposite.

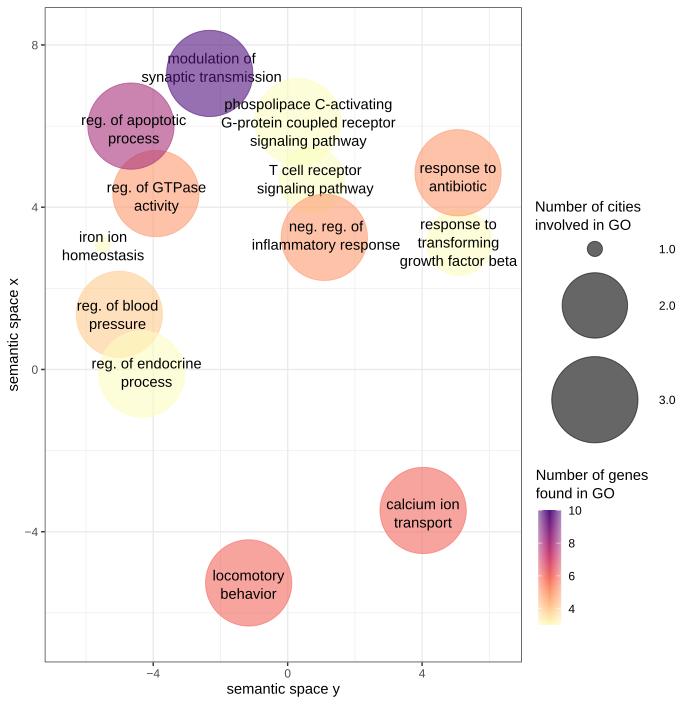

**Figure S5:** Semantic space of enriched GO terms from genes associated with urbanization-linked DMRs. Semantic space coordinates have been calculated based on similarity of GO term word composition using REVIGO. The color gradient indicates the number of genes involved in each GO term and circle size indicates the number of city contributing to each category.

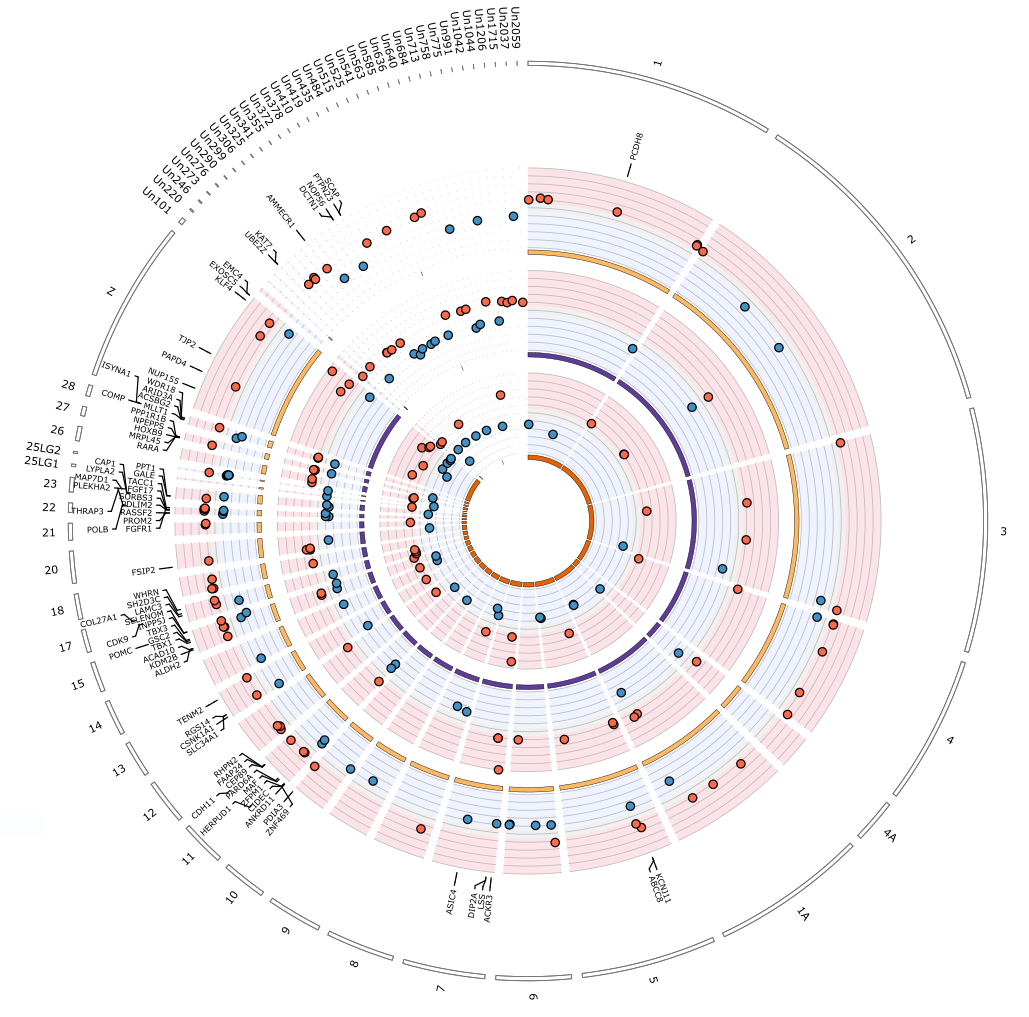
**Figure S6:** Circos plot of differentially methylated regions (DMRs) identified between females and males great tits in and near Barcelona, Montpellier and Warsaw (from inner to outer circles). Red points show hypermethylated regions in female great tits relatively to males, and blue points show hypomethylated regions. For graphical clarity, only a subset of genes are represented: genes associated with the 10% most extreme DMR. Names of the genes found within 5 kb of the represented DMRs are given.

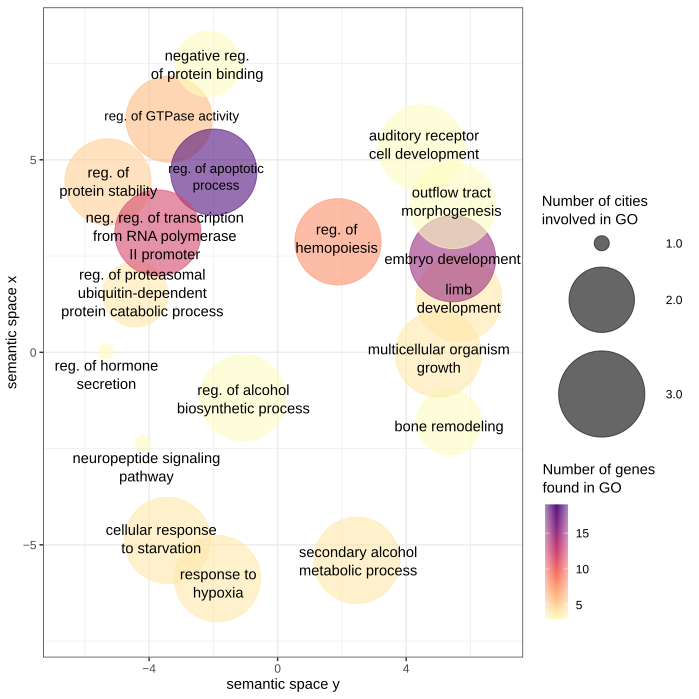

**Figure S7:** Semantic space of enriched GO terms from genes associated with sex-linked DMRs. Semantic space coordinates are calculated based on similarity of GO term word composition using REVIGO. The color gradient indicates the number of genes involved in each GO term and circle size indicates the number of city contributing to each category.

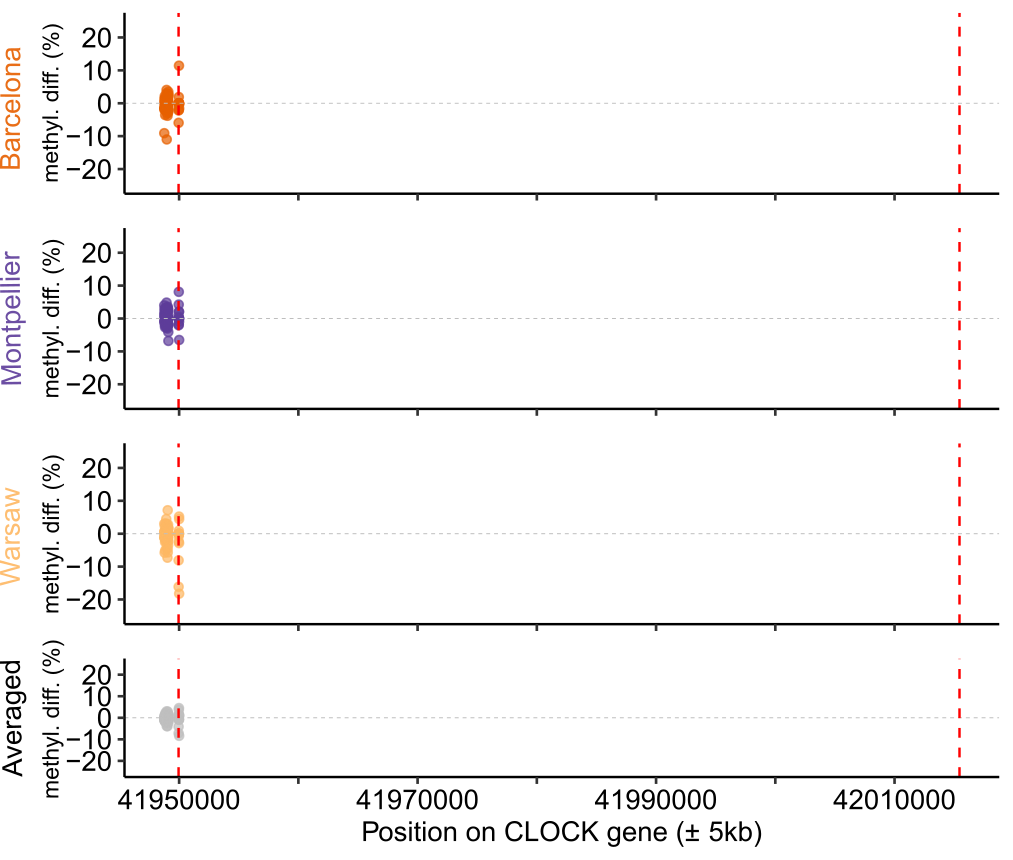

**Figure S8:** Mean methylation difference per base along the CLOCK gene. Vertical dashed lines materialised start and end of the gene.

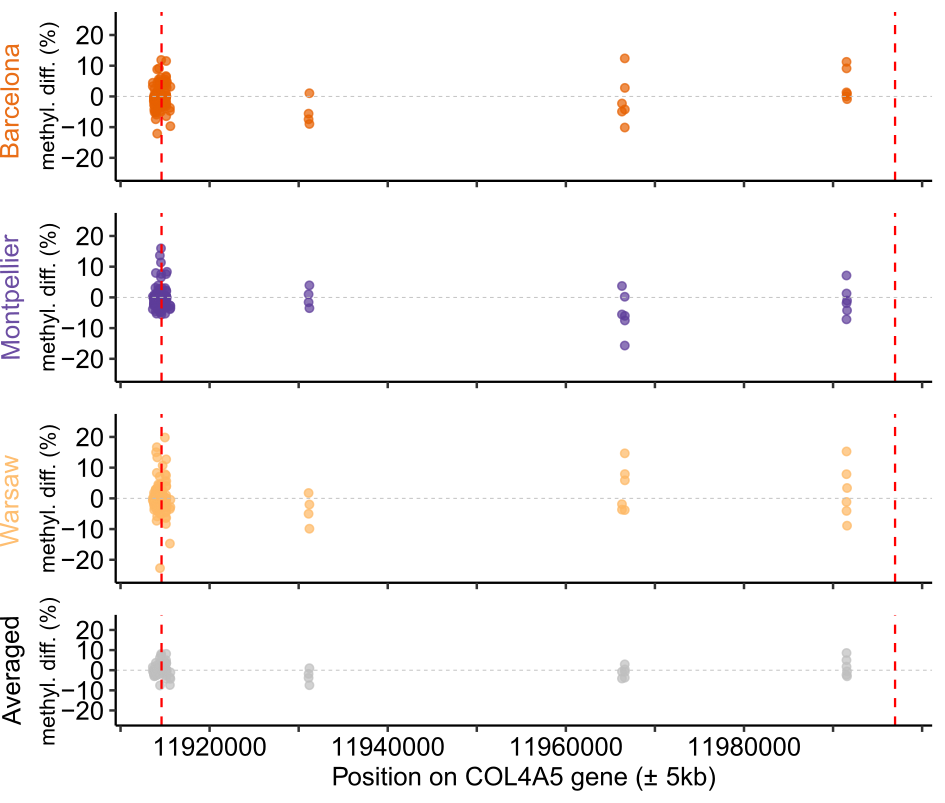

**Figure S9:** Mean methylation difference per base along the COL4A5 gene. Vertical dashed lines materialised start and end of the gene.

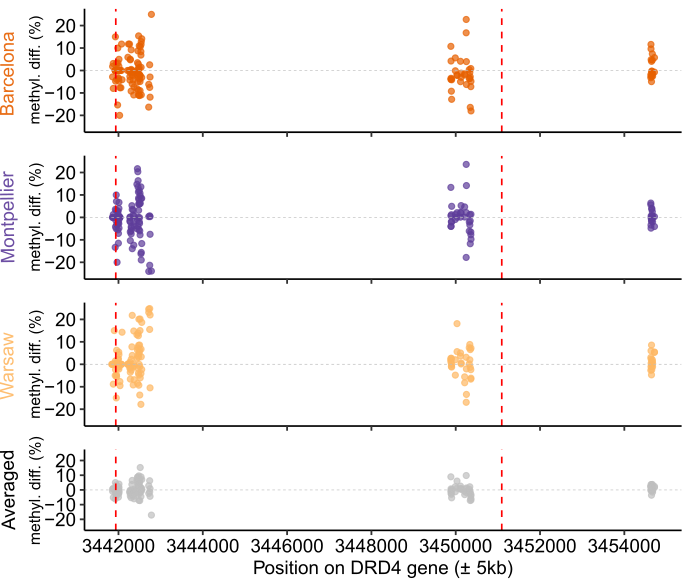

**Figure S10:** Mean methylation difference per base along the DRD4 gene. Vertical dashed lines materialised start and end of the gene.

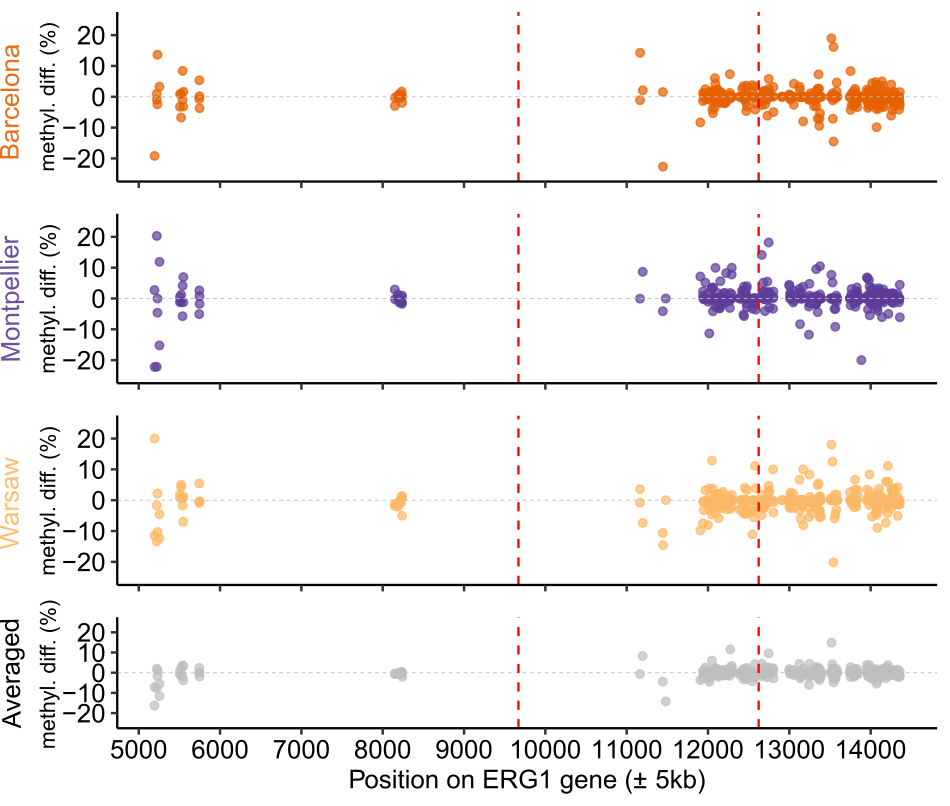

**Figure S11:** Mean methylation difference per base along the ERG1 gene. Vertical dashed lines materialised start and end of the gene.

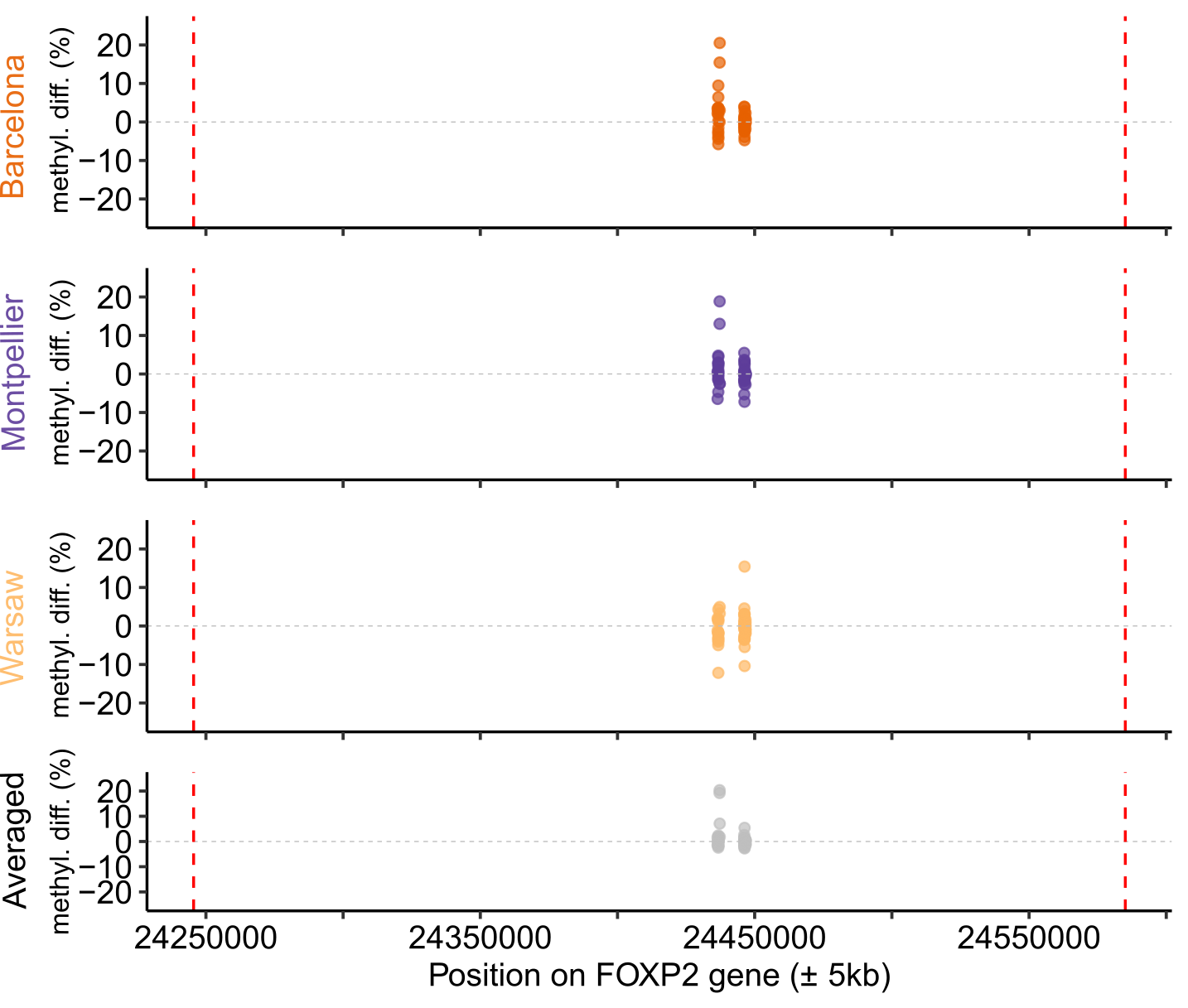

**Figure S12:** Mean methylation difference per base along the FOXP2 gene. Vertical dashed lines materialised start and end of the gene.

**SUPPLEMENTARY ANALYSES**

**Differentially Methylated Regions (DMRs) between Females and Males**

A total of 238 DMR were found between sexes: 58 in Barcelona, 81 in Montpellier and 99 in Warsaw (Figure S6). Warsaw presented significantly more hyper than hypomethylated DMR (χ^2^=5.878, P= 0.015), but it was not the case for Barcelona (χ^2^=0, P=1) nor Montpellier (χ^2^=1.25, P=0.264). DMR were distributed on 29 chromosomes and 35 unplaced scaffolds.

In total, 181 DMR were located on genes or in 5kb upstream/downstream regions around genes. A gene ontology enrichment analysis performed on the pooled list of genes associated with DMR revealed significantly enriched modules associated with metabolism, development and hormonal function (see details Figure S4B and S6, and Table S11).

**Differentially Methylated Regions between associated with urbanization in females and males**

To explore the potential sex dependent effects of urbanization on methylation patterns, we searched for DMRs between forest and urban habitats in females and males separately. The numbers of significant DMRs per sex per location are reported in Table S12.

In females, 101 DMRs (7.9%) were shared between 2 or more locations and 45 (3.5%) showed the same direction of variation in all cities.

Similarly, in males 134 DMRs (9.7%) were shared by multiple locations, and 58 (4.18%) showed the same direction of variation.

Within locations, sexes showed shared urban-linked DMRs, see Table S12.
